# Brawn before bite in endemic Asian eutherian mammals after the end-Cretaceous extinction

**DOI:** 10.1101/2025.09.24.678280

**Authors:** Z. Jack Tseng, Qian Li, Suyin Ting

## Abstract

The first 10 million years (Myr) following the Cretaceous-Paleogene (K-Pg) mass extinction marked a period of global greenhouse conditions and dramatic rise of placental mammals. Because ∼80% of known terrestrial sections capturing post-K-Pg mammal recovery come from North America, a substantial knowledge gap exists in the tempo and mode of recovery in Asia, where only 3% of global sites are located and most contain species found nowhere else. We show that isolated Paleocene eutherian assemblages from China (1) exhibited high mean tooth size and disparity early in the Paleocene, (2) shifted in their dental shape in parallel with regional and global environmental changes later in the Paleocene, and (3) achieved maximum dental shape-performance covariation near the end of the first 10 Myr post-K-Pg. This ‘brawn before bite’ transformation, coupled with prolonged dental shape versus performance variability, favors a scenario whereby many living orders of eutherian mammals were borne out of phenotypically and functionally plastic ancestral assemblages, including those in tropical south China, during the Paleocene.

## Main Text

The Cretaceous-Paleogene (K-Pg) mass extinctions accelerated the formation of modern-day global biota. In particular, the more than 6,000 species of living placental mammals trace their origins to the diversification of major orders around or after the K-Pg boundary ^1–3^. Different hypotheses (for example: early rise, suppression, and late rise) about the timing of their adaptive radiation have been proposed ^1,4,5^. Regardless of the diversification scenarios favored by the competing explanations, the Paleocene epoch (66-56 Myr ago) has been highlighted as among the most critical time interval for establishing the macroevolutionary parameters of the placental adaptive radiation ^6,7^ (but see ^5^). However, substantial asymmetry exists in the quantity and quality of terrestrial fossil records that document the first 10 Myr of the ‘Age of Mammals’, with ∼80% of known terrestrial K-Pg boundary sections worldwide occurring in North America ^8^. Earliest Paleocene mammals are not known from Europe ^9,10^, rendering detailed reconstructions of post-K-Pg recovery in that continent difficult. The post-K-Pg fossil assemblages from Asia have hitherto not been considered in analyses of post-K-Pg recovery dynamics ^11^.

The North American fossil record suggests that post-K-Pg eutherian taxonomic diversity recovery was relatively rapid; most occurred within the first ∼0.3 to 1 Myr of the Paleocene epoch ^12–17^. This initial eutherian diversification was driven by archaic groups (i.e., stem placental/eutherian lineages and extinct placental subgroups), followed by the first appearance of many modern orders during two peak hyperthermal events in the past 66 Myr, at the end-Paleocene and early Eocene climatic maxima, respectively ^18,19^.

Distinct from these taxonomic recovery patterns, high selectivity of mammalian ecomorphological extinction across the K-Pg boundary indicates a primary productivity filter at the start of the Cenozoic Era (66 Myr ago to the present) ^20^. Additionally, the first 10 Myr of mammalian brain evolution after the K-Pg was marked by niche partitioning along a size gradient; endocranial traits reflecting more complex sensory processing did not appear until the Eocene (56 Myr ago). This phenomenon has been termed the ‘brawn before brains’ hypothesis ^21^. A similar pattern in initial size-driven diversification followed by expansion of ecomorphological disparity is also observed in mammal jaws, indicating a general dynamic across phenotypic systems ^22^. Whether this mode of evolution, whereby size disparity increase precedes other ecomorphological traits, should be understood as a global phenomenon during the post-K-Pg placental radiation remains untested, in large part because no such analyses have centered on non-North American continental fossil mammal records.

A major challenge with expanding analyses of post K-Pg recovery to Paleocene mammal assemblages elsewhere in the world is the stratigraphically limited nature of early Cenozoic sequences that produce fossil mammals. In Asia, Paleocene localities in China represent the best studied to date ^11^. From the earliest Paleocene, highly regional and endemic faunas are known from a handful of sedimentary basins (Fig. S1A). Among the recorded faunal elements, only the archaic placental clades Anagalida and Pantodonta are consistently sampled across the major subdivisions of the Paleocene ^11^. Additional complications with ecomorphological analysis of these stem eutherians include the uncertainty in their dietary ecology, having diverged prior to the crown radiation, and uncertainty in phylogenetic positions of Paleocene taxa ^7^; thus, they are beyond the reach of conventional phylogenetic bracketing approaches to dietary reconstruction. Phenomic analysis of the placental radiation supports insectivory as the ancestral diet of the hypothetical placental ancestor, but uncertainty in the post K-Pg availability of insects and plants in some regions leave some doubt as to the accuracy and scope of this ancestral state reconstruction ^1^. Herein we treat the archaic Paleocene taxa in our analyses as having uncharacterized diets rather than categorizing them as insectivores, herbivores, or carnivores.

We investigated the timing of ecomorphological diversification by developing and leveraging the largest dataset to date of Paleocene Asian eutherian assemblages. Our analyses focused on eutherians from three of the most fossiliferous and biogeographically isolated Paleocene sedimentary sequences in paleotropical Asia: The Nanxiong, Qianshan, and Chijiang Basins in present-day south China ^23–27^ (Fig. S1). We generated a new phenotypic dataset of 200 Asian Paleocene eutherian teeth using high-resolution microcomputed tomography and laser scanning, capturing 37 species endemic to low-latitude east Asia and which are brought to bear on K-Pg recovery dynamics for the first time (Data S1-S2). Teeth are among the most well-preserved parts of fossil mammals, and the fact that they interface directly with the environment through mastication makes them suitable elements for studying potential ecology-morphology linkages. We used dental topographical traits as indicators of ecomorphological diversity ^28^ and examined temporal shifts in tooth crown complexity, curvature, and height. We additionally assessed the association of topographical traits with tooth crown mechanical performance in terms of deformation resistance using topographic and simulation analyses. Our dataset spans the Paleocene, the first 10 Myr of the Cenozoic, enabling us to test the hypothesis that dental topography and tooth puncturing and shearing performance linkages showed delayed niche expansion relative to mean body size and size disparity during this initial period of post-K-Pg eutherian recovery in Asia.

### Dental traits paralleled Paleocene global and regional environmental conditions

We posit that dental topographic trait variability in Paleocene eutherians in south China tracked global and regional climatic changes despite stasis in high-level taxonomic composition over the course of the first 10 Myr after the K-Pg transition (Fig. 1). Dental height and sharpness variability were low in the beginning and end of the time interval, with a peak in the middle Paleocene. This pattern is observed both when dentitions are considered as pooled samples through time, and particularly driven by the lower dentition (Fig. S5; note that upper teeth display the opposite pattern). In contrast, elevated low-level taxonomic turnover of genera and species within the Paleocene indicates cladogenetic shifts, rather than anagenetic adaptation, underpin this dental topographic evolution (Data S1)^11^. These findings suggest that in addition to its impact on crown group eutherians, the K-Pg extinctions and subsequent climatic fluctuations played a role in filtering out archaic eutherians with lower speciation rates in favor of rapidly speciating taxa ^29^.

**Fig. 1.**
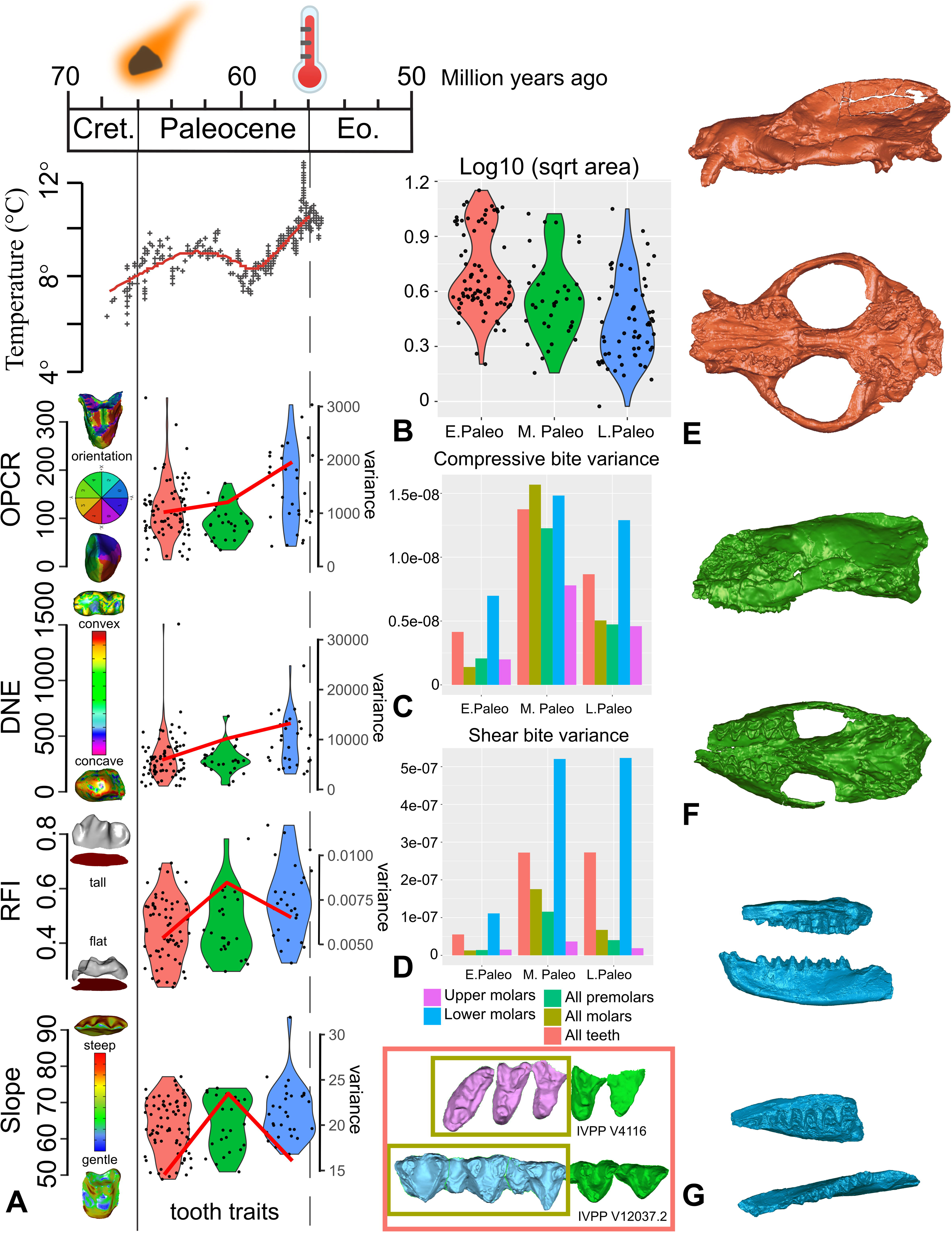
Temperature and fossil eutherian mammal dental trait shifts during the first 10 Myr of the Cenozoic. (A) Dental topographic trait values (boxplots) and mean variance (red curves) during the first 10 Myr of the Cenozoic, signifying the time after the K-Pg mass extinctions and before the Paleocene-Eocene hyperthermal event. Global temperature curve based on Zachos et al. ^61^. Dental traits measured include crown complexity (OPCR, orientation patch count rotated), curvature (DNE, Dirichlet normal energy), height (RFI, relief index), and slope. (B) Eutherian tooth size distributions represented by log 10 square root tooth area, in units of log10 millimeters. (C) Variance of compressive bite performance based on tooth crown finite element simulations, in units of squared Joules. (D) Variance of shear bite performance based on tooth crown finite element simulations, in units of squared Joules. Examples of endemic Asian fossil specimens analyzed: (E) Lateral and ventral views of early Paleocene Chinese endemic pantodont (CEP) *Bemalambda nanhsiungensis* IVPP (Institute of Vertebrate Paleontology and Paleoanthropology, Chinese Academy of Sciences) V4116. (F) Lateral and ventral views of middle Paleocene CEP *Harpyodus decorus* IVPP 5035.1. (G) Lateral and occlusal views of late Paleocene CEP *Guichilambda zhaii* IVPP V12037.2 (dentary) and V12037.3 (maxillary fragment). Firey asteroid symbols indicate the end-Cretaceous asteroid impact in the Yucatán Peninsula; thermometer symbols indicate the Paleocene-Eocene hyperthermal event. Subdivisions of the Paleocene approximately correspond to the Shanghuan, Nongshanian, and Gashatan Asian Land Mammal Ages, respectively (see supplemental text for competing age boundary scenarios).

By contrast, we found no support for significant shifts in dental topographic trait mean values from the early to the middle Paleocene for the majority of analytical iterations of clade and dental data partitions (Figs 1A, S4; Table 1; Data S7). However, in most analyses we observed a significant shift in at least one dental topographic metric from the middle to the late Paleocene (Table 1; Data S7). The larger-bodied Chinese endemic pantodont (an ungulate-like archaic placental group; “CEP”) mammals tend to increase dental complexity (OPCR, orientation patch count rotated) and curvature (DNE, Dirichlet normal energy) whereas smaller, non-pantodonts (Chinese arctostylopids and anagalids; rodent-like archaic placental groups) in the dataset exhibited no significant dental trait shifts from the middle to late Paleocene (Fig. S4; Data S7). The transformation by CEPs in complexity and curvature indices relates to the capacity of teeth to resist wear and coincides with temperature and aridity increase towards the end-Paleocene thermal maximum event (Figs. 1-2). Multiple geological and geochemical proxies suggest that paleoclimate in and around the Nanxiong Basin K-Pg section in south China reflects a latitudinally much broader global tropical zone during the Paleocene ^30^ relative to present-day Earth, as well as rapid shifts between more versus less humid intervals during the first 10 Myr of the Cenozoic ^31^. Results from feldspar-quartz (F:Q) ratios, clay mineral composition analysis, diffuse reflectance spectroscopy (DRS), stable carbon isotope analysis, and total organic carbon (TOC) analysis all support this regional paleoclimate profile ^31,32^. Local climate reconstructions for the latest Cretaceous indicate relatively warm and dry intervals, followed by warm and humid climates during the earlier Paleocene, then a return to less humid but still warm conditions in the later Paleocene ^32^. Chinese endemic pantodont dentitions tracked shifts in this paleoenvironmental progression.

**Fig. 2.**
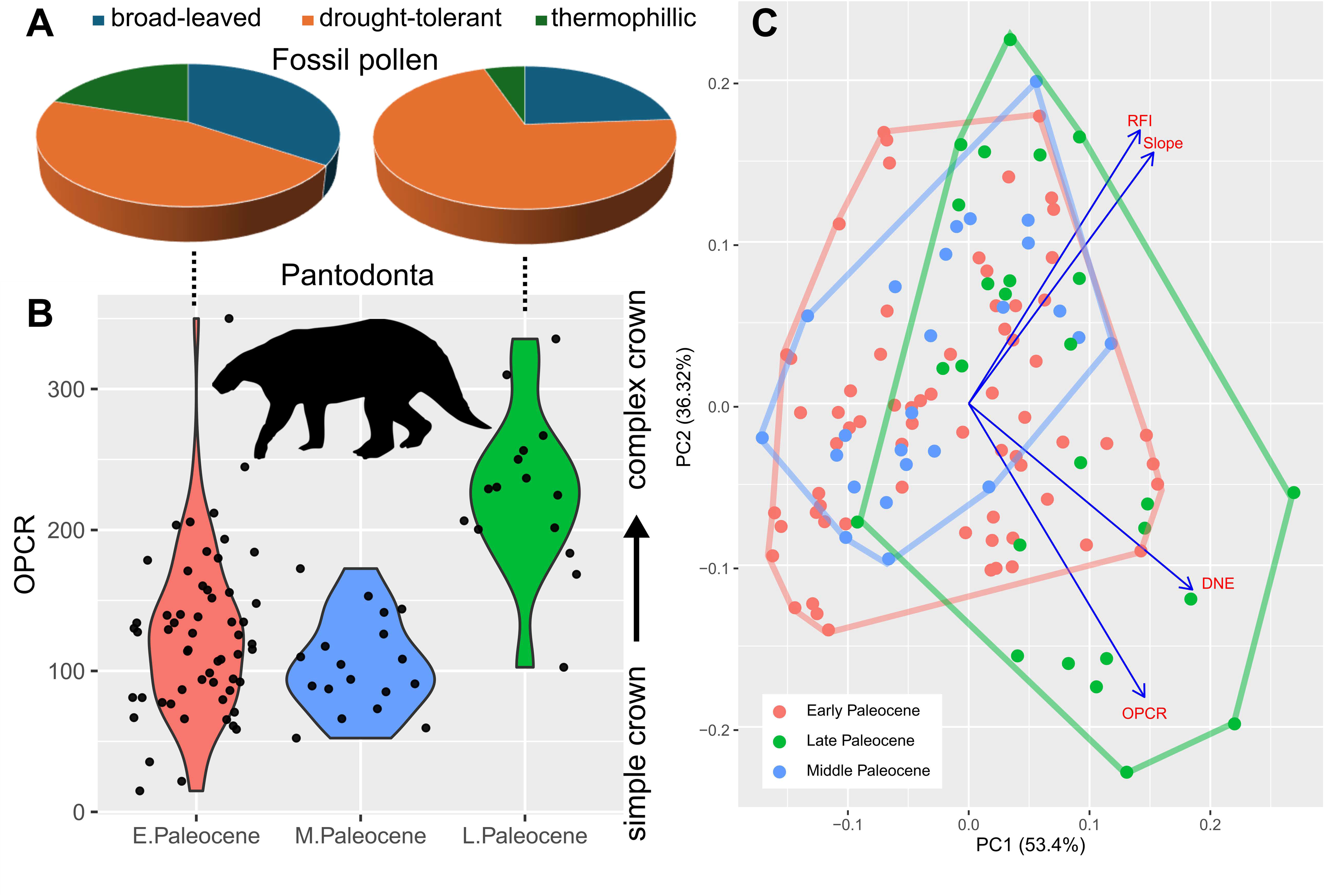
Association of paleopalynological data from the Nanxiong Basin, south China, and late Paleocene niche expansion in endemic Asian fossil eutherians. (A) Proportion of environmental humidity indicator taxa from early versus late Paleocene paleobotanical localities, respectively, in the Nanxiong Basin; data based on ^35,36^(Data S12). (B) Boxplots of dental complexity (OPCR, orientation patch count rotated) in the Chinese endemic pantodont (CEP) data partition across the three Paleocene time intervals examined. Note the concomitant increase in CEP tooth complexity (OPCR) and increased proportion of drought-tolerant plant species in the Nanxiong Basin during the late Paleocene. (C) Principal component morphospace of all tooth data analyzed; convex hulls delineate overall morphospace occupation during each time interval. Eigenvectors of the four dental topographic traits are indicated in blue. Late Paleocene shift and expansion in dental topographic morphospace is statistically significant at the *p* = 0.05 level (Table 1). Pantodont silhouette by S. Traver from phylopic.org.

**Table 1.**
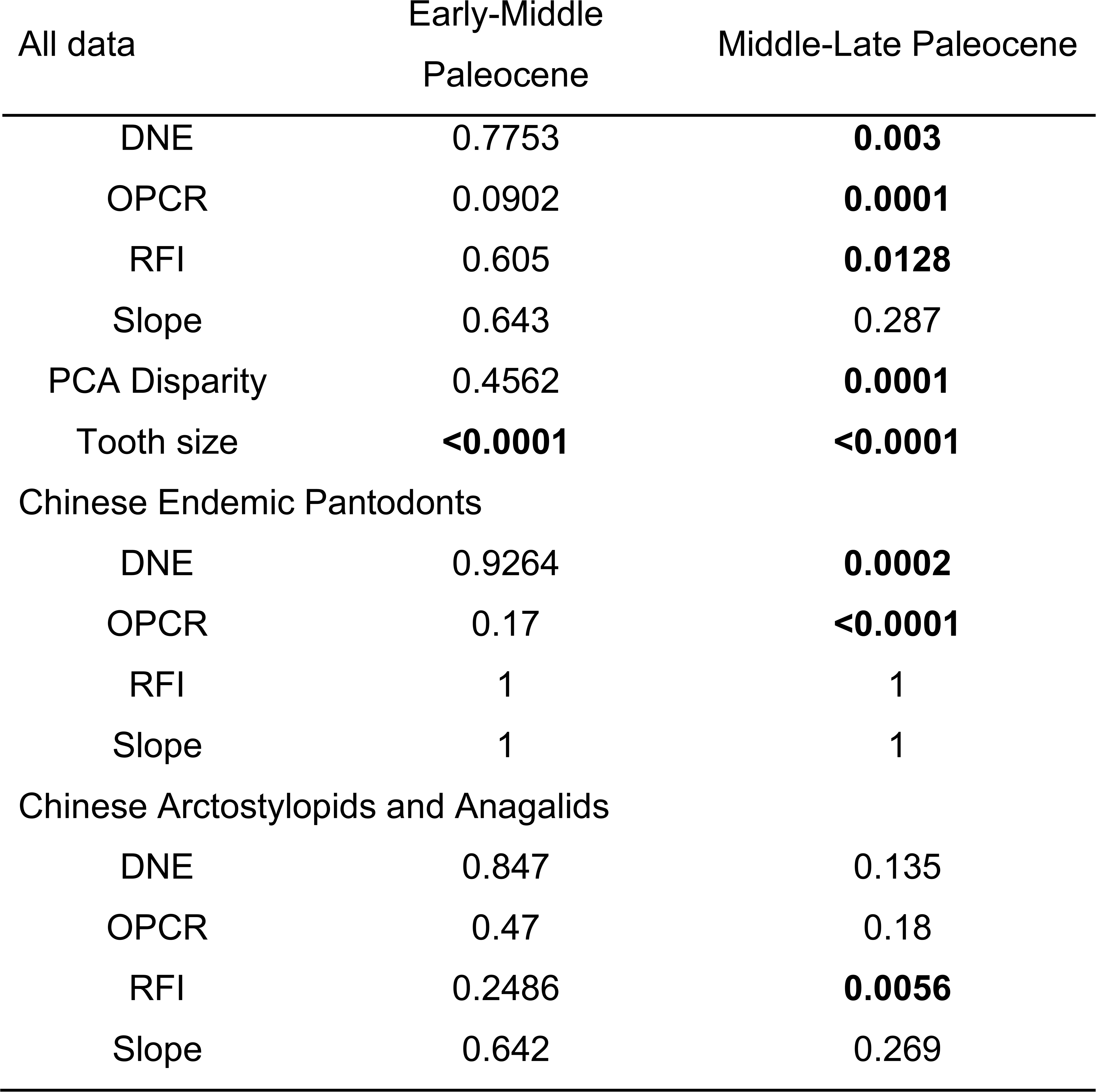
Pairwise t test of dental topographic trait and disparity differences across adjacent time bins. Dental topographic trait differences are assessed across time intervals in all-data, Chinese endemic pantodont, and non-pantodont partitions. Dental trait disparity was estimated based on all principal component axes using the outputs of PCA. Tooth size variance differences were calculated from tooth area or square root of tooth area in all-data and no-outlier partitions to assess effect of outliers on statistical significance (see Data S6 for details). Bolded font indicates *p* values < 0.05.

The overall increase in dental complexity and curvature also coincided with an increase in drought-tolerant flora in south China (Fig. 2) and specifically with paleoenvironmental reconstructions in the Nanxiong Basin ^33^. Palynological evidence suggests that in addition to a predominance of broad-leaved, deciduous plants mixed with a smaller percentage of conifers than in fossil localities further north ^34^, south China also recorded drought resistant taxa such as the maidenhair fern *Pterisisporites* in the early Paleocene Shanghu formation ^35^ and again in late Paleocene samples ^36^. Expanded comparisons across time and space suggest fossil pollen taxa that are indicators of seasonal aridity increased from 20.3% of late Paleocene pollen samples to 34.3% of early Eocene pollen samples ^36^. These shifts appeared to have stabilized by the end of the Eocene, when taxonomically modern floras became established in Asia ^37^. In addition to tracking Paleocene temperature trends (Fig. 1), Asian eutherian dental topography (OPCR and DNE in particular) also mimicked trends in the marine realm, where global planktonic foraminifera records demonstrate a similar pattern in species richness curve, as well as with global δ^13^C ^38,39^ and south Pacific CO_2_ levels ^40^. These broader associations underscore the inference that endemic Paleocene eutherians in south China comprised a dynamic assemblage shifting their dental morphology in step with regional and global environmental changes during the first 10 Myr after the K-Pg mass extinctions.

We detected a shift and expansion of eutherian dental morphospace in the late Paleocene (Fig. 2). An overall shift towards increased dental topographic trait magnitudes in late Paleocene samples is driven mainly by CEPs (Table 1), despite the fact that they constitute a minority of the overall data (41% of teeth) as well as late Paleocene partition (21% of teeth). Additionally, dental metric disparity is significantly higher in the late Paleocene partition than in the two preceding time bins (Table 1). This pattern is driven both by increased dental curvature and complexity (DNE & OPCR; 78-100% support in per-tooth analyses, Fig. S5) in larger-bodied CEPs and crown height (RFI) in smaller-bodied non-pantodonts, in addition to differences in disparity among the clades (Data S7, Fig. S5; bootstrapped variance for DNE, OPCR, and RFI are at least twice as large in non-pantodonts compared to CEPs, and bootstrapped range lengths returned similar patterns). This suggests that an expansion of dental disparity in the late Paleocene occurred across the size spectrum of endemic Asian eutherians. Over the same time interval examined, body (tooth) size disparity and mean were higher in the early Paleocene than in subsequent time intervals (Fig. S8, Table S3; also supported by premolar 4 and upper molar partition analyses), indicating that substantial increases in the disparity of dental complexity, curvature, and height lagged behind tooth size during the Paleocene. Dog-sized CEPs such as *Bemalambda* reached sizes not seen in late Cretaceous mammals from China such as *Zhangolestes* and *Kryptobaatar*, which are shrew- to gopher-sized ^41^. This suggests a ‘brawn before bite’ pattern in endemic Asian eutherians, partially mirroring the endocranial and jaw functional morphology patterns identified in their North American and European counterparts ^21,22^. These findings raise the possibility that an initial size-driven post-K-Pg recovery followed by ecomorphological radiation was a global phenomenon, even as regional tectonic events such as the initial collision of the Indian subcontinent with Asia and Deccan Traps volcanism influenced local mammal evolution ^42,43^.

### Topography-performance covariation underlies eutherian dental shifts

Within an overall pattern of increasing covariation between dental topographic traits and bite performance traits across the Paleocene time intervals (Fig. 3; Table S1), topographic versus performance trait variability shifted both from early to middle and from middle to late Paleocene time bins (Figs S3-S4). Early Paleocene to middle Paleocene DTA-FEA intra- and inter-correlations remained stable (Fig. 3B), but DTA-FEA inter-correlations strengthened while intra-DTA correlations weakened in the Middle to Late Paleocene transition (Fig. 3C). This transition pattern is in part due to the divergence of shape-performance linkages in CEP versus non-pantodont eutherians, and is particularly driven by the upper and lower first molars (Figs. S9-S11). Dental topography variability tracked bite performance variability in non-pantodonts through time, peaking in the middle Paleocene (Fig S4D). By contrast, both intra- and inter-partition comparisons of topography and performance trends in CEPs showed low correlations through time (Fig. S4G). The coexistence of distinct topography-performance relationships in each time and taxon partition while overall covariation between the two trait groups increases between time bins is consistent with form-function decoupling ^44^. Complex form-function linkages generally promote evolutionary redundancy and can enhance optimization of phenotypic traits when selective trade-offs are present ^44–46^. The presence of functional redundancies underlies the high levels of dental topographic variability in Asian Paleocene eutherians. Alternatively, varying degrees of independence between the two performance traits and dental topographic traits analyzed could allow the two aspects of the dentition to evolve in a decoupled manner (Fig. 3).

**Fig. 3.**
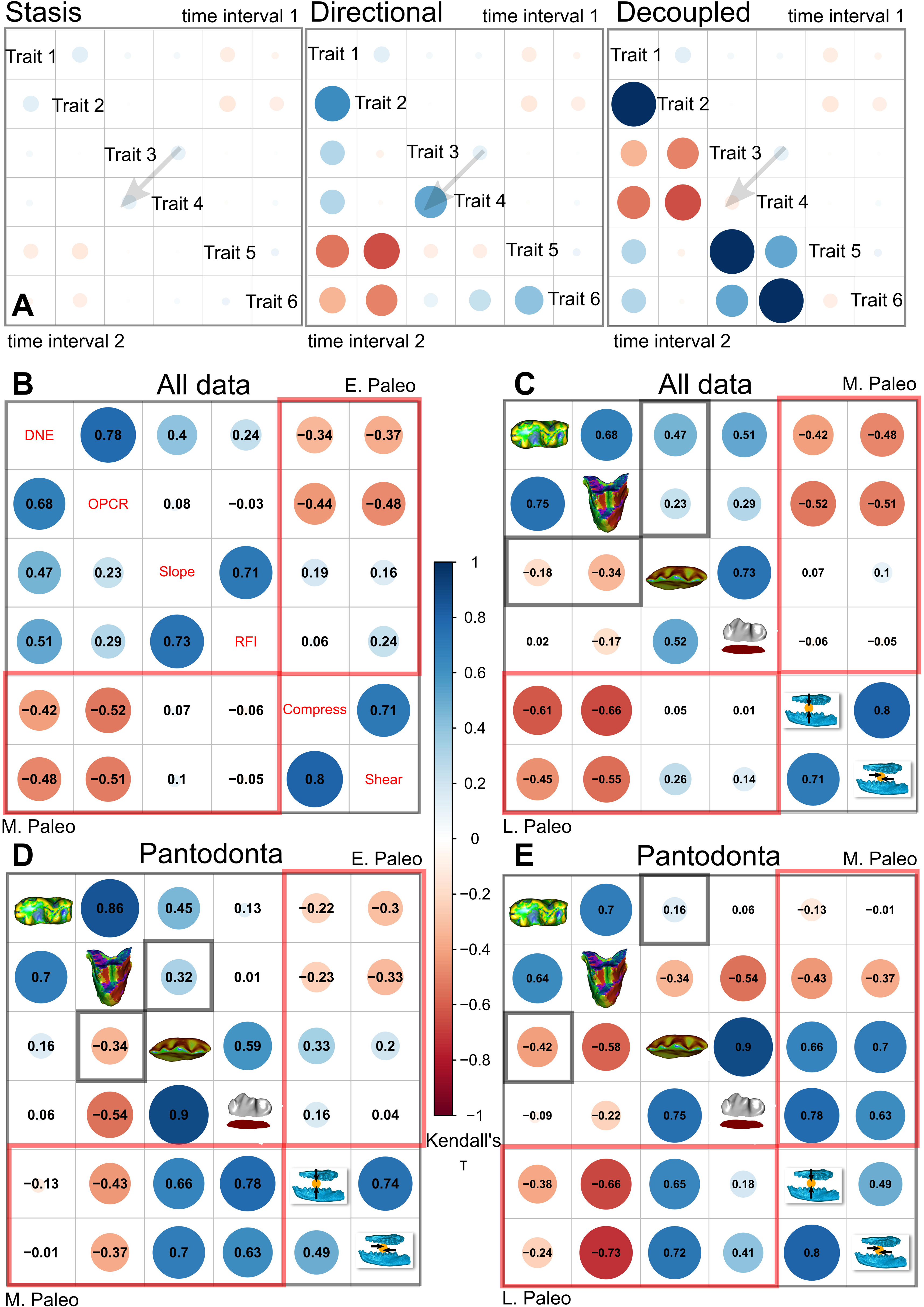
Correlation plots of dental topographic and bite performance traits in endemic Asian Paleocene eitherians. (A) Hypothetical correlation scenarios used to interpret stasis, directional, verses decoupled change through time in specimen data. (B) Pairwise ranked correlation coefficient estimated using Kendall’s τ between early and middle Paleocene dental topographic and performance traits in the main dataset. (C) Correlation between middle and late Paleocene traits in the main dataset. (D) Correlation between early and middle Paleocene traits in the Chines endemic pantodont (CEP) data partition. (E) Correlation between middle and late Paleocene traits in the CEP data partition. Topography-performance correlations are marked in red boxes. Decoupled/reversed trait correlations are marked in gray boxes.

### South Asia as a Paleogene ‘Garden of Eden’ for eutherian mammals?

Among the most consequential implications of accurately interpreting post-K-Pg mammalian recovery dynamics in Asia is the ability to reconstruct the evolutionary conditions during the biogeographic origination of modern placental orders. Current knowledge and an overwhelming majority of data about the first post-dinosaur mammal ecological communities center on North American localities ^12,47^, and secondarily from European records ^48^. Yet, the fossil records of those continents pinpoint a high turnover of mammal faunas at the Paleocene-Eocene boundary, driven by the Cretaceous/Paleocene origination of major modern mammal clades elsewhere and subsequent dispersal of those lineages to the European and North American continents, respectively. At least five living orders (Primates, Rodentia, Lagomorpha, Perissodactyla, Eulipotyphla), representing over half of mammalian species richness today, trace their evolutionary origins to Asia ^18,49–51^. There is also mounting evidence that other organisms, including fish and plant lineages, followed a similar biogeographic pattern ^52^.

The abrupt appearance of early representatives of modern mammal lineages in North America has been formally operationalized by the ‘East of Eden’ model, wherein Asia is the ‘Garden of Eden’ and a biodiversity pump of living orders of eutherian mammals ^53^ and other organisms. Biogeographic modeling analysis of modern mammalian diversity also strongly identifies tropical south Asia, including the geographic regions occupied today by southern China, India, and southeast Asian countries, as the cradle of mammalian diversification since the Paleocene ^54^. As geographically proximate faunal assemblages to the epicenter of the ‘Garden of Eden’ hypothesis, the endemic Paleocene eutherians of the Asian paleotropics analyzed in this study exhibited a high degree of dental topographic variability while strengthening topography-performance covariation during a period of biogeographic isolation and regional and global climate changes. Such flexibility in dental form-function linkage permits ‘mix and match’ trait combinations rather than evolutionary transformation as a single unit, potentially enhancing the evolvability of feeding ecological traits as new environmental conditions arose ^56^. This finding favors the scenario whereby many living orders of placental mammals were borne out of phenotypically and functionally plastic ancestral assemblages, including those that lived in south China, during the Paleocene ^55^.

We further hypothesize from these findings that episodes of fluctuating global warming during the first 10 Myr of the Cenozoic promoted the evolution of ‘all-purpose’ mammalian dentitions rather than those with specialized functions (Fig. 3). This prolonged dental topographic variability tracks extended post-K-Pg floral recovery times ^36,57^, and suggests a ‘brawn before brains’ mode of placental mammal evolution may in part represent a series of correlated evolutionary shifts in sync with the steadily increasing complexity of Paleocene primary producers ^20,37^. That a global signal would be detected in the splendidly isolated south Asian Paleocene assemblage is notable, and indicates that effects of climate forcing on post-K-Pg mammal recovery may be ubiquitous, as has been predicted for Earth’s ongoing rapid environmental shifts ^58^. In response, Paleocene mammal clades in south China were relatively larger early on and increased covariation between dental topography and bite performance later, all the while maintaining high levels of variability in dental complexity and convexity (Fig. 1). These preconditions may have set the stage for the subsequent taxonomic turnover between archaic and crown lineages during the Paleogene to Neogene modernization of mammal communities ^19,26^. As a primary interface against shifting food resources and the environment, the eutherian dentition was poised to play an outsized role in the explosive diversification following the K-Pg extinctions, with the masticatory complex having already completed the macroevolutionary transition to a mechanically stiff but phenotypically flexible jaw during the Mesozoic Era ^59,60^. The end-Cretaceous extinctions and the climatic volatility that followed then set in motion the ecological release that enabled mammals to explore dental form-function to a far fuller extent in 66 Myr of the Cenozoic Era than during the preceding 150 Myr under the reign of dinosaurs. This new hypothesis of ‘brawn before bite’ in tooth size, topographic, and performance evolution further refines the placental success story.

## Acknowledgments

We thank I. Ruiz for assisting with generating 3D tooth models from CT data; X. Tseng for assisting with data collection; P. Holroyd for assisting with locating specimen casts in the collections of the University of California Museum of Paleontology; J. Liu for helpful discussion and comments on earlier versions of the study. The High-Resolution X-ray Computed Tomography Laboratory, Institute of Vertebrate Paleontology and Paleoanthropology, Chinese Academy of Sciences, provided assistance with the CT scanning. Z. Luo provided earnest and constructive feedback on an earlier version of the study. Editor S. Rasmann and two anonymous reviewers provided highly constructive feedback on the study.

## Author contributions

Conceptualization: All authors

Methodology: ZJT.

Formal analyses: ZJT.

Investigation: All authors.

Results acquisition: ZJT.

Raw data acquisition (surface and CT scan data): ZJT, QL.

Writing, original draft: ZJT.

Writing, review, and editing: All authors.

Visualization: ZJT, QL.

## Resource availability

- Requests for further information and resources should be directed to and will be fulfilled by the lead contact, Jack Tseng.
- Code associated with analyses is included with this article.
- Raw data are available in the supplementary materials.
- All original fossil specimens are accessioned in the Institute of Vertebrate Paleontology and Paleoanthropology, Chinese Academy of Sciences.

## Supplemental information

Document S1. Figures S1–S11, Tables S1–S3, and supplemental references

Data S1

Data S2

Data S3

Data S4

Data S5

Data S6

Data S7

Data S8

Data S9

Data S10

## METHOD DETAILS

We focused sampling on three clades: Chinese endemic Pantodonta, Chinese Arctostylopida, and Anagalida. Additional data were collected on other clades (e.g., Chinese Tillodontia) opportunistically when well-preserved specimens were available (Data S1). These three main clades together represent >50% of the species found in Paleocene faunas across China ^11^. They also individually represent the most diverse clades across the three Asian Land Mammal Ages of the Paleocene: Shanghuan, Nongshanian, and Gashatan. By contrast, these clades are reduced to <25% of species diversity in Eocene assemblages in China, finally disappearing altogether in the late Oligocene. Therefore, we take the three clades to be representative of eutherian assemblage dynamics during the Paleocene time interval in China.

Historically, Paleocene fossil mammal faunas in China have been defined based on biostratigraphic criteria and supplemented by magnetostratigraphic correlations where available. Here we follow the assessment of ^62^ in generally correlating the Shanghuan Asian Land Mammal Age (ALMA) with the Puercan and Torrejonian North American Land Mammal Ages (NALMAs), the Nongshanian ALMA with early to middle Tiffanian NALMAs, and the Gashantan ALMA with late Tiffanian and Clarkforkian NALMAs, respectively (see ^11^ for an alternative interpretation). Additionally, the Shanghuan and Nongshanian boundary was interpreted by ^25^ to be at the top of Chron C27N, which is dated at 60.920 Myr ago in the Geomagnetic Polarity Time Scale ^63^ but indicated as closer to ∼62.3 Myr ago in ^62^. Similarly, the Nongshanian-Gashatan boundary has been variably defined at 59.24 Myr ago ^62^ or the base of Chron C26N, which is 57.911 Myr ago ^63^. In contrast, the end of the Gashatan coincides with the Paleocene-Eocene boundary at 56 Myr ago and has been consistently defined as such ^62^. Given these existing uncertainties, we use the terms ‘early Paleocene’, ‘middle Paleocene’, and ‘late Paleocene’ to refer generally to the Shanghuan, Nongshanian, and Gashatan ALMAs, respectively, in this study.

We analyzed 200 individual teeth from 48 specimens, representing 37 species (Data S1-S2). The teeth represent 2 upper first premolars, 4 upper second premolars, 15 upper third premolars, 19 upper fourth premolars, 22 upper first molars, 23 upper second molars, 16 upper third molars, 3 lower first premolars, 5 lower second premolars, 13 lower third premolars, 17 lower fourth premolars, 20 lower first molars, 22 lower second molars, and 19 lower third molars. These tooth positions were selected from a broader examination of ∼300 individual teeth from 72 specimens. We vetted the specimens and excluded 99 tooth positions (∼33% of teeth initially chosen for possible inclusion) from our analyses because they either (1) were partially or completely broken at the crown, (2) were in an advanced stage of attritional wear where no cusps could be identified, or (3) possessed a combination of the two aforementioned conditions. The assigned geologic age of the studied specimens spans the entire Paleocene, binned into early (n = 91 teeth), middle (n = 46 teeth), and late Paleocene (n = 63 teeth) data partitions. To maximize sample size while minimizing disturbance to more delicate specimens, a combination of original specimens and previously produced high-fidelity specimen casts were used in our sampling of dental crown morphology.

Given the rarity of Paleocene fossil material from China, we combined data from different tooth positions into three pooled samples, one for each of the time intervals examined (early, middle, late Paleocene). We treated the pooled samples as representative of the range of dental topographic features and bite performance traits available to the eutherian taxa under study. In this way, the variance estimates are interpreted as measures of the morphological and performance heterogeneity present in each time interval dataset. To further tease out the possibility of specific tooth positions driving the overall trends observed in the pooled samples, we also performed the DTA, FEA, DTA-FEA correlation, and tooth size through-time analyses using per-tooth data partitions.

### Tooth shape estimates using Dental Topographic Analysis

We digitized dental crown morphology via microCT either using a GE v|tome|x m 300/180 kV micro-computed-tomography system (GE Measurement & Control Solutions, Wuntsdorf, Germany), housed at the Institute of Vertebrate Paleontology and Paleoanthropology (IVPP), Chinese Academy of Sciences (CAS), or a GE Phoenix Nanotom M in the Functional Anatomy and Vertebrate Evolution Laboratory, University of California, Berkeley. Projection images (1,000 to 1,500 images depending on specimen size) were acquired with an isotropic voxel size of 10 to 40 μm, at an energy range of 120-150 kV, current of 100-150 μA. Additionally, surface 3D scans (generated using IVPP’s Artec Space Spider 3D scanner, with a precision of 0.05 mm and resolution of 0.1 mm) were used for larger specimens to efficiently obtain surface morphology data. The enamel caps (the crown portion of each tooth above the enamel-dentine junction) were extracted from the specimen models using Geomagic Wrap v2021; all crown surface holes (generated from scanning imperfections or digitally removed sediment) were patched, the models were remeshed with triangles that have aspect ratios (base:height or edge-edge) of <10 and decimated to ∼10,000 triangles. Finally, we standardized the spatial orientation of all tooth crowns before exporting them as .ply files for dental topographic analysis (DTA). Individual tooth crown (enamel cap) models have the Z axis normal to the occlusal plane, the X axis parallel to the mesial-distal axis, and the Y axis parallel to the labial-lingual axis.

Exported meshes were further vetted and prepared for DTA within the R programming environment. We used functions implemented in the *molaR* R package ^64^ for all of the steps described next. First, all .ply files are further cleaned using ‘molarR_Clean’ to remove floating points/vertices and any triangular faces with zero area. Then, the batch function ‘molarR_Batch’ was used to calculate four dental topographic metrics: DNE, OPCR, Slope, and RFI.

Dirichlet Normal Energy (DNE) is a measure of occlusal sharpness, using a quantification of surface energy in tooth crowns relative to a gently curving or flat mesh surface. Steep, high, and/or shearing cusps tend to produce higher DNE values in DTA, whereas bulbous cusps tend to produce lower DNE values ^65,66^. DNE has been used to successfully distinguish folivores, omnivores, and frugivores in euarchontan (primates, colugos, tree shrews) mammals ^65,67^.

Orientation Patch Count Rotated (OPCR) is a measure of tooth crown complexity, using quantification of distinct patches on the crown face that possess unique orientations. Teeth with larger number of cusps and crenulations tend to produce higher OPCR values, whereas teeth with fewer cusps and simpler ridges tend to produce lower OPCR values. This metric has been used effectively to distinguish convergently evolved carnivore versus herbivore species across multiple mammalian orders ^68^. DeMers and Hunter ^69^ suggest that OPCR does not currently provide a reliable approximation for the degree of adaptation to herbivory, and that any interpretation of ecomorphological differences should be made within a taxon-specific context. Given the narrow set of clades targeted in our dataset, we interpret increasing OPCR as an indication of increased functional capability to process vegetation. A related measure, slope, quantifies the mean tilt of the tooth surface relative to the occlusal plane. Teeth with lower and more gently curving cusps tend to have lower slopes, whereas teeth with sharp and/or abruptly ridged crests tend to have higher slopes.

Relief Index (RFI) is a measure of the relative height and complexity of the tooth crown, using the values of the 3D surface and 2D ‘footprint’ areas of the tooth crown. RFI has been shown to be effective in distinguishing frugivores from insect and leaf specialists in euarchontan mammals ^65^. Furthermore, RFI is also able to distinguish carnivoran species at different trophic levels and dietary breadths ^66^.

Overall, we use these DTA traits as indicators of ecomorphological capacity, but do not link them explicitly to dietary categories. The craniodental morphology of archaic placental clades in general have not been demonstrated to share the same structure-function linkages as crown mammals, so aforementioned linkages between DTA and dietary ecology from studies of extant species only serve as evidence that DTA is a potentially useful ecomorphological proxy. We do not directly apply those DTA-diet relationships to the Paleocene fossil eutherian dataset.

All tooth crown models are provided as ply files and deposited in FigShare (10.6084/m9.figshare.28611854).

### Tooth performance estimates using Finite Element Analysis

The common inference that dental morphology can reflect dietary adaptations makes an underlying connection between the two traits through biomechanical performance; the mechanical performance imbued by a particular morphological configuration accomplishes one or more food acquisition and/or processing tasks. Such mechanical performance links can be tested using experimental and simulation approaches on both theoretical grounds and actual tooth shapes ^70^. Here we applied a simulation approach to estimate two general performance traits commonly interpreted for mammalian tribosphenic teeth: puncturing/compressing and shearing/grinding ^71^. The ancestral therian tribosphenic stroke, or the chewing movement underlying mammals with unfused mandibular symphysis and tribosphenic molars, has been reconstructed to involve significant components of (1) long-axis rotation in each hemimandible and (2) mortar-and-pestle grinding between upper and lower cusps ^72^. These movements are associated with the ‘lock-and-key’ occlusion of a stereotypical mammalian tribosphenic dentition, which exhibit tall crest-like cusps in the trigonid and lower basin-like valleys in the talonid of lower molars that articulate with their corresponding upper teeth (Fig. S3).

We used finite element analysis (FEA) to estimate the work efficiency of different tooth crowns subjected to compressive/puncturing and shearing/grinding forces, the two major masticatory actions of tribosphenic teeth ^71,73,74^. Adequate compressive and puncturing forces applied through individual or combined action of cusps are needed to overcome the fracture strength of brittle and tough food such as seeds and invertebrate exoskeletons in order for individuals to access soft and/or more nutritious tissues within. Sufficient shearing and grinding forces applied through surfaces or cusp edges are necessary to further break down food boli into smaller pieces to aid in the digestive process. The combination of these two major functions of tribosphenic teeth is thought to be a foundational adaptation that enabled the mammalian radiation ^71,72,75^. We focused on the dental enamel portion of the tooth crown in our biting simulations because it is the most mineralized vertebrate tissue and typically the best-preserved component of fossil teeth. The fracture mechanics of mammalian dental enamel is thought to be intimately related to dietary function ^76^, including the evolution of different enamel thickness in different feeding ecomorphologies ^28^. It is important to note that because the dental data available for this study included not only CT scans, but also surface scans and specimen casts, enamel thickness was not quantifiable for most specimens in our dataset (see below for a description of enamel thickness scaling procedure applied in the simulations).

The two masticatory scenarios tested, compressing/puncturing versus shearing/grinding, have previously been shown to be broadly connected to tooth topology. Taller cusps (approximated in our study by Relief Index and partially by Slope) and more convex crowns (approximated in our study by DNE) tend to exhibit higher strain in finite element models of hypothetical tooth shapes ^77^. However, there is by no means a consensus on the presence of a tight form-function linkage across all tooth types and/or clades. Whereas a dental topography to food mechanical property linkage can be detected in some extant carnivores ^78,79^, bats ^80^, and primate-like tooth models ^81^, a form-function relationship between dental topography and tooth/food mechanical properties is absent in other model systems ^67,82^. Therefore, we incorporated biomechanical simulations as an explicit test of the mechanical performance associations implied by the DTA-flora correlations established in the dental topography and paleoenvironmental portion of our analyses. The main question addressed by these simulations is not necessarily whether a form-function linkage exists in the Paleocene eutherian dataset, but whether there is a consistent relationship between DTA and FEA traits across the three time bins of the Paleocene. If such consistent linkages are observed, it would suggest a stasis in dental topography and compressive/shearing performance through the first 10 million years of the Cenozoic in our dataset; alternatively, any significant changes in the topography-performance relationship would indicate substantial evolutionary change through the same time period.

We imported the same tooth crown models generated for DTA into the Strand7 finite element software (Strand7 Pty Ltd, Australia) to create finite element models of the individual teeth. The tooth meshes are composed of two-dimensional, three-noded triangular elements ranging between 9,000-10,000 elements, with assigned thickness parameter (see below) so they behave similarly to single-layer 3-D elements. A convergence test was done on two of the specimens (IVPP V5228, V5231) and indicated that the <10,000 three-noded triangular plate elements (tri3) returned simulation output values with less than 10% deviation from higher resolution meshes of the same models (Fig. S3). Therefore, we deemed the default resolution of the tooth meshes used in DTA to be sufficient for finite element simulations. Furthermore, because the dataset contained digital models built from both CT and surface scan data, not all specimens have enamel thickness information. Although increased enamel thickness has been associated with durophagy in mammals, there appears to be taxon-specific patterns of enamel thickness to tooth size and dietary ecology, in addition to extensive variation in intra-tooth enamel thickness distributions ^83–85^. In absence of information on enamel thickness differences among the taxa studied, we standardized the thickness of the tooth crown models isometrically to be 10% of the total surface area of a given tooth model. As such, the simulation results should be treated as enamel cap shape-derived performance traits, rather than those based on a fully parameterized enamel model incorporating localized thickness and inter-specific allometric differences.

We scaled applied forces on the teeth, simulating compressive or shearing loads, to be a value equivalent to total model surface area (but in units of Newtons instead of mm^2^). Compressive loads were represented on each tooth by a single nodal force vector directed towards the base of the crown along the height axis of each tooth, on the tallest cusp or cuspid (which is typically the protocone for upper molars and protoconid for lower molars, or the main cusp in premolars). This configuration simulated a compressive load directed into the tallest cusp (towards the root) on a given tooth from a food item (Fig S3), as reconstructed for the mammalian tribosphenic bite during a crushing or puncturing movement ^71,72^. Shearing loads were represented on each tooth by two nodal force vectors, one on each of the two tallest cusps or cuspids; the two nodal forces are of equal magnitude but opposing directions, perpendicular to the long axis of the tooth and in the occlusal plane. This configuration portrays a shearing motion along the short axis of the tooth (Fig. S3), and simulates tooth-to-tooth and tooth-to-food contact during the ‘mortar and pestle’ grinding rotation movement reconstructed for the ancestral mammal tribosphenic bite ^72^. This force magnitude standardization procedure effectively generates identical force to area ratios across the models, ensuring that the magnitudes of mechanical stress placed on the model are the same across models of different sizes; furthermore, we adjusted the strain energy values collected for each tooth model simulation using the volume and input force ratio-based correction equation provided in ^86^. Given our objective to assess the association between relative performance traits and DTA metrics, with the latter being size-free variables, only the relative magnitudes of bite performance, rather than absolute values, were collected from the performance analyses.

After each tooth crown model was defined using the criteria outlined above, homogenous material properties were assigned. All models were defined as plate models (with thickness scaled to surface area) with a Young’s (elastic) modulus of 80 GPa ^87^ and Poisson Ratio of 0.3. Next, each tooth was constrained from translation and rotation using four nodal constraints distributed in the four corners of each tooth, respectively (Fig. S3).

All bite scenarios were solved using the linear static solver implemented in Strand7. A total number of 400 analyses were performed (one compressive bite and one shearing bite simulation for each of the 200 tooth specimens in the dataset). We extracted two traits from the models: compressive bite strain energy and shear bite strain energy. Strain energy is defined as the area under a stress-strain curve of an object under load; operationally it measures the amount of work done by the load to deform the object (analogous to experimental work-to-fracture measurements)^88^. An object that is more resistant to deformation would have lower strain energy than an object that easily deforms under load. The use of work-to-fracture measures to assess tooth performance is consistent with the understanding of mammalian tooth enamel as a fracture-prone, yet fracture-resistant, biological tissue ^89^. In such a framework, fracture resistance is expected to be directly related to the amount of force an individual is able to exert through the tooth crown during mastication, and by extension the hardest or toughest food item that can be processed without catastrophic damage to the tooth crown itself. Fracture resistance (as approximated by strain energy in this study) is also likely to be a stronger target of selection in mammals compared to those of other toothed vertebrates, given the former’s diphyodonty (having only two sets of tooth instead of continuous dental replacement) and thus necessity to prolong usable tooth lifespan^83^. Therefore, we used strain energy values under compressive and shear bite simulation scenarios as a proxy for the effectiveness of each tooth crown model at resisting deformation from the respective biting forces. A tooth that is more resistant (i.e., have lower strain energy values) from compressive forces would be able to more effectively crush/puncture harder food items, and a tooth more resistant from shear forces would be able to cut/grind tougher food items. We modify Irschick et al.’s^90^ definition of whole-organism performance and define bite performance in the context of this study as the capacity of individual tooth models to resist simulated compressive and shearing forces, which represent ecologically relevant factors that influence masticatory efficacy.

All finite element models are provided as NASTRAN/text files and deposited in FigShare (10.6084/m9.figshare.28611854).

### Sensitivity validation of original versus cast specimen models

The tooth dataset developed in this study contains a mixture of CT scans of original and casts of fossil specimens. In order to assess the potential discrepancies in DTA and FEA trait values collected from cast versus original specimen derived models, either due to phenotypic details not captured in cast specimens or deterioration of epoxy or resin-based casts over time, we randomly sampled two specimens for which both original and cast based models were analyzed. Results from DTA and FEA of the CT-derived models of lower first molars from IVPP V5228 (*Altilambda pactus*) and V5231 (*A. tenuis*) were compared to those conducted on digital models built from CT scans of casts of those specimens.

We used the ‘n-point’ and ‘global’ alignment functions in Geomagic Wrap to first align the models using arbitrary homologous landmarks visually identified on both tooth models and then the automatic global alignment function to maximize overlap between the two models in 3D coordinate space. The ‘deviation’ function in Geomagic Wrap was then used to calculate summary statistics for linear deviations orthogonal to the surface of the original model. The V5228 Lm1 model showed a maximum deviation of 0.58 mm (5.8% of crown height) and average deviation of 0.04-0.09 mm (<1% of crown height), with a standard deviation of 0.11 mm (1% of crown height). The V5231 Lm1 model showed a maximum deviation of 0.38 mm (6% of crown height) and average of 0.03-0.04 mm (<1% of crown height), with a standard deviation of 0.06 mm (1% of crown height)(Fig. S3).

### Quantification of uncertainty ranges for bootstrap sampling

We then subjected the cast-based models to DTA and FEA. In both cases, the DTA values between cast and specimen models are the most different for DNE (30-40% difference), followed by OPCR (18-32% difference), then RFI (2-13% difference), and finally Slope (1-5% difference) (Data S4). Also in both cases, adjusted compressive SE values differed by 14-41% and adjusted shear values differed by 20% (Data S3). Based on these specimen-cast validation tests, we set a conservative +/- 40% uncertainty range for all DTA and FEA values obtained from all cast-based tooth models. DTA and FEA trait mean and variance estimates used for downstream statistical analyses were then calculated using a bootstrap sampling scheme where 1,000 replicates of the DTA and FEA datasets were passed through the statistical tests (see Quantification and statistical analysis section, below). Overall, the early and middle Paleocene time bins contain around 50% cast data, whereas the late Paleocene time bin contains 66% cast data. If data uncertainty from original versus specimen models present a major signal, we would expect the late Paleocene time interval to always show higher variance given an abundance of cast data in that time bin. We did not observe any consistent trends of high variance in the late Paleocene in the 1,000 bootstrap samples, suggesting that the results are not significantly biased by data quality differences between cast and original specimen derived models (Fig. S4).

### Time bin duration correction and tooth position ratios

According to Ni et al. [6], the Shanghuan Asian Land Mammal Age (ALMA) faunas in Guangdong and Anhui range from 66-61.6 Ma (or 4.4 Myr in duration), the Nongshanian ALMA faunas in Guangdong, Jiangxi, and Anhui range from 61.6-59.2 Ma (or 2.4 Myr in duration), and the Gashatan ALMA faunas in Anhui range from 59.2 to 56 Ma (or 3.2 Myr in duration). Given the different durations represented in each of the three time bins used in our analyses, we corrected all variance estimates by dividing the variance calculated in each of the bootstrapped samples by the time duration of the respective time bin. In this manner, the variance values reported in the study represent per-million-year values.

We used pie chart analysis to verify that the proportions of tooth positions in each time bin are not substantially different from each other (Fig. S1). However, we caution that time bin comparisons of individual tooth positions are unlikely to be statistically robust because of small sample sizes. We only use aggregates of dental positions (all teeth, molars, or premolars) in our data partition analyses and interpretation.

## QUANTIFICATION AND STATISTICAL ANALYSIS

We used the following features of the dental topography and performance data as the basis for assessing faunal assemblage trait shifts through the Paleocene: trait mean, variance, trait-to-trait correlation, and partition-to-partition correlation. We used ANOVA (Analysis of Variance) and pairwise t test to compare trait means, F test to compare trait variance, linear regression analysis and Kendall’s τ to compare trait-to-trait correlation, and two-block partial least squares (2B-PLS) analysis to compare partition-to-partition correlation. There are a total of six traits forming two main data partitions: topographic partition (DNE, OPCR, Slope, RFI) and performance partition (Compressive SE, and Shear SE). To assess the sensitivity of the results to subsets of the data, all of the trait comparisons were done iteratively using the total dataset, by time bin (early, middle, late Paleocene), taxon (Chinese endemic pantodonts versus non-pantodonts), and/or by dental position (all teeth, molar teeth, premolar teeth). All statistical tests were performed on 1,000 bootstrap resampled datasets that were parameterized using results of sensitivity tests on original versus cast specimen models, finite element mesh convergence tests, and time bin duration comparisons (see previous section). In addition, we quantified tooth size using two measures: total 2D surface area and square root of surface area. The R script for all statistical analyses and plots are included in Data S9.

### Statistical resampling and tests

We used bootstrap resampled variance of dental topographic metrics and dental performance variables as a measure of tooth form-function disparity. 1,000 replicates of the 200-specimen DTA-FEA trait datasets were generated by sampling each DTA and FEA trait value from a uniform distribution defined using a +/-40% range of the original model-derived values. The bootstrap resampling was done with replacement, so it was possible for a given DTA/FEA trait value in a sampled tooth model to be repeated in a different replicate, but very unlikely for entire datasets to share similar values with another replicate. We used this resampling scheme to account for the uncertainty in trait estimates introduced by the totality of modeling, specimen preservation, cast versus original, and other potential sources of uncertainty in trait values. For each replicate, variance was calculated for each metric, time interval, for Chinese endemic pantodont (CEP) versus non-pantodont, and molar versus premolar data partitions. To assess shifts through time, statistical differences between the variance of each dental topographic metric sample in adjacent time intervals (A, early Paleocene; B, middle Paleocene; C, late Paleocene) were evaluated using F tests. A null hypothesis of a variance ratio of 1 between pairwise comparisons was tested using the var.test() function in R ^91^. The outputs of the variance tests on a given bootstrap sample were then compiled for all 1,000 replicates, and the overall variance mean and variance test output used as the statistical basis for detecting significant differences between time, taxon, and tooth data partitions. Unless indicated otherwise, all tests described below were also done using the 1,000 bootstrap samples.

We evaluated differences in central tendency (mean value) of dental topographic metrics and dental performance variables through time using analysis of variance (ANOVA) and pairwise t tests. Each dental topographic metric was tested against the three time intervals defined above, using the aov() function in R. For statistically significant (at the *p* < 0.05 level) results, we additionally evaluated the pairwise intervals that contribute significant differences in mean dental topographic values. We assessed pairwise differences using pairwise *t* tests implemented with a Holm correction for multiple comparisons. The *t* test was conducted using pairwise.t.test() in R.

We constructed a tooth morphospace using all four dental topographic metrics analyzed by principal components (PC) analysis. The first and second PC axes were chosen to visualize the two-dimensional morphospace. Additionally, we quantified the degree of morphological disparity and statistical differences in disparity between adjacent time interval data partitions. All PC scores generated from the PCA were included in the disparity analysis. The prcomp() function in R was used for PCA, and the morphol.disparity() function implemented in the Geomorph R package ^92^ was used for morphological disparity significance tests. Input data for the PCA were from the original dataset values, not from the bootstrap samples.

Linear regression analysis between individual dental topographic metric (DNE, OPCR, Slope, RFI) and dental performance metric (compressive bite strain energy, shear bite strain energy) was performed to quantify the correlation between dental form and function. Adjusted *R*^2^ and *p* values generated from the lm() function in R were used to evaluate the goodness of fit and statistical significance of the form-function relationships. The distribution of *R^2^* values and their corresponding *p* values are reported in Fig. S6.

In addition to pairwise form-function linear regression analyses, we also evaluated the degree of correlation between the DTA and FEA data blocks as a means to measure covariation between dental topography and deformation resistance. We used two-block partial least squares analysis^93^ coupled with bootstrap resampling to generate distributions of correlation coefficients (*r-pls*) for each of the three time bins, and then tested for significant differences in the magnitude of DTA-FEA correlation between adjacent time bins using Welch’s two-sample t tests on the correlation coefficient distributions. DTA-FEA correlation differences at *p* < 0.05 are interpreted as a significant shift in the degree of covariation of the two traits from one time bin to the next time bin.

All R scripts used in the analyses are included as supplementary files (Data S8, S9).

