## Supplemental Information for "Brawn before bite in endemic Asian eutherian mammals after the end-Cretaceous extinction"

Supplemental Document

Field locality information

The extensive development and conversion of outcrops in the Nanxiong Basin into agricultural fields has created a ‘race against time’ to document and study this critical area to understand placental evolution. Clyde et al. [1] estimated that 20 out of 54 early and middle Paleocene fossil sites in the Nanxiong Basin have already been destroyed by infrastructural and housing developments since their original discovery nearly three quarters of a century prior. Our re-survey of the key fossil sites during a trip there in 2023 suggests that the majority of the sites, except for the K-Pg boundary locality, which is protected by municipal historic landmark designation, are now inaccessible or completely obliterated. This reality means that the mammalian fossil samples analyzed in this study offer an ever more important and rare window into earliest mammal life during the ‘Age of Mammals.’ Our analyses are based on the most complete earlier Paleocene materials currently recovered from the Asian continent.

Isotope analyses show that average δ^13^C values returned to pre K-Pg levels from a short-duration 2 ppm (part per mil) decrease about 1 Myr into the Paleocene in the Nanxiong Basin, congruent with global patterns of rapid mammal taxonomic recovery [1]. Although the majority of Paleocene fossiliferous localities are within the classic ‘red beds’ of south China, interbedding with limestone-like layers may indicate localized cycles of dry and humid intervals [2]. Fossil pollen in the Nanxiong Basin indicates warm climates throughout the Paleocene time interval; however, a notable shift in higher percentages of ferns and gymnosperms and lower percentage of angiosperms is observed in the late Paleocene [3], indicating local recovery of gymnosperms and ferns (Table S2).

The antipodal location of the Nanxiong Basin from the Yucatan Peninsula makes the geochemical identification of a K-Pg boundary layer difficult. Arguments for the precise location of the K-Pg boundary layer has been made on the basis of iridium enrichment in dinosaur egg shells [4] and with total mercury content that correlates with known timing of Deccan Traps volcanism [5]. Regardless of the precise stratigraphic level of the K-Pg boundary in the Nanxiong Basin, the specimens we included in our analyses have all been collected in higher stratigraphic levels that are biostratigraphically unambiguous as Paleocene sequences [3]. Planned fieldwork at and around the putative K-Pg boundary sections will sample micromammal fossils towards the objective of constructing a refined biostratigraphic framework for high-resolution analysis of the first million years of post-K-Pg mammal recovery. At this time, high-resolution data are not available to test the precise rate at which Paleocene Asian mammal taxonomic disparity and size disparity recovered/stabilized.

Evolutionary preconditions for the post-K-Pg placental radiation

Ting et al. [6] described an *in situ* faunal turnover event in southern China between the early and middle Paleocene and a gentle decrease in endemic taxa. Such a faunal change did not amount to shifting dental topographic mean values, although it is correlated with increased variability in all topographic traits (Fig. 1). Similar inferences of ecological stasis were made by Clyde et al. [7]. The replacement of some archaic taxa with more cosmopolitan ones is consistent with an evolutionary ‘training ground’ phenomenon where taxa preadapted to local environmental conditions were at a selective advantage to disperse into new geographic regions when global climate changed [8]. Hooker [9] makes the observation that late Paleocene North American and European mammal assemblages responded to the late Paleocene thermal maximum by increasing browsing herbivory and terrestrial taxa; this increase is at the expense of arboreal taxa in the European samples analyzed in that study. The observed DTA shifts in our dataset are consistent with this trend (Fig. 2). In the final Paleocene time bin, immediately prior to the faunal turnover that coincided with the Holarctic mammalian dispersal event and the PETM (Paleocene-Eocene Thermal Maximum), increased seasonal aridity in southern China (based on pollen composition) correlated with increased disparity and maxima of dental topographic metrics (Fig. 2).

The majority of research into post-K-Pg mammal recovery come from North American sites, which represent ~80% of known terrestrial K-Pg boundary sections [10]. By contrast, Asian K-Pg sites represent only ~3% of K-Pg boundary sections. There are no Paleocene fossil mammal assemblages known from the Indian subcontinent [11]. Europe is similarly problematic in terms of both the paucity of well-dated K-Pg sites as well as correlated or thick stratigraphic sections that permit research into post-K-Pg dynamics [12]. Additionally, the oldest well-sampled Cenozoic mammal fauna in Europe is late early to middle Paleocene in age. This absence of earliest Paleocene data renders the study of mammalian evolutionary dynamics from the K-Pg to end-Paleocene hyperthermal event in Europe difficult [13]. Thus, no dental form-function comparisons are possible between the Asian and European post-K-Pg mammal recovery patterns for the entire Paleocene time interval.

Bayesian analyses of Paleogene mammal faunas suggest that a zoogeographic barrier between the Arabian Peninsula and the rest of Asia was present and prevented faunal interchange until the end-Oligocene [11]. Thus, the corridors of dispersal and interchange for the southern Chinese Paleocene mammals at the end-Paleocene were constrained to the north, towards the Mongolian Plateau, or to the south towards India. The absence of Paleocene Indian fossil mammals prevents a characterization of the extent to which the ecomorphological patterns described in this study also applies more broadly to fossil mammals further to the south. Based on the establishment of a land connection between Asia and India during the Paleogene, similarity in ecomorphological patterns might be expected [14]. By contrast, the well-known Paleocene-Eocene stratigraphic sequence and fossil record on the Mongolian Plateau clearly document the northward expansion of early Paleocene southern Asian taxa and the increased faunal similarity with other continents [15] during time intervals immediately following those analyzed in this study.

Absence of latitudinal gradients in Paleocene mammal taxonomic richness despite modern level temperature gradients across western interior North America suggests potential ecological instability in that region around the time of the mammalian dispersal to North America [16]. However, other analyses suggest that arrival of Eurasian immigrants to North America at the Paleocene-Eocene Thermal Maximum did not drastically alter functional diversity in the local communities [17]. Establishment of latitudinal floral gradients and the high mammal endemism in east Asia during the same time period [18] may have allowed ecological stability to establish earlier in Asia compared to North America. The stability of a large landmass in Asia for much of the Mesozoic and Cenozoic likely also contributed to the development of climate-adapted faunas in south Asia leading up to the PETM and expansion of suitable warm habitats for the southern taxa [19]. Data presented here suggest that adaptations of endemic east Asian mammals to global and regional environmental changes in the critical 10 Myr period immediately following the K-Pg extinctions can be understood as priming the evolutionary pump for the origin, radiation, and dispersal of modern mammal orders.

Summary of levels of topography-performance associations

(1) Correlation plot patterns (Fig. 3): We performed correlation plot comparisons on combined DTA and FEA data, using Kendall’s τ as a measure of the strength of correlation. Contingency tables show that DNE/OPCR to compress SE/shear SE correlations increased from early to middle then late Paleocene (Fig. 3). Compared to the steady increase in integration shown by two-block PLS analyses (see below), this suggests that dental complexity and convexity were disproportionately driving the overall integration patterns, whereas cusp height and sharpness were very weakly associated to compressive and shear strain energy values.

We additionally conducted correlation analyses across time using data partitions representing individual tooth positions (Figs. S9-S11) to verify the overall trend observed using the total dataset (Fig. 3). Upper M1 patterns generally reflect the trend recovered from analysis of the overall dataset, but M2 and M3 results display inconsistent DTA-FEA correlations, possibly due to small sample sizes. Lower molar patterns generally replicate those recovered in the overall analyses, but lower M1 and M2 signals appear to be stronger than those for lower M3. Finally, low sample sizes make premolar-specific correlations unstable, with general pattern showing EP-MP strengthening then MP-LP stasis or weakening.

(2) Per-time variance patterns (Figs. S3-S5): The overall dataset showed steady increase in DNE and OPCR variance, differing from the middle Paleocene peak pattern seen in the RFI, Slope, Compressive SE, and Shear SE variances. The non-pantodont partition shows a more consistent pattern between DTA and FEA variances, with all traits showing a middle Paleocene spike. Lastly, the CEP data partition shows even more mosaic patterns than the overall data partition. CEP DNE, OPCR, and Slope show variance increases over time, whereas Slope and compressive SE exhibit a middle Paleocene spike. Shear SE shows a steady decline in variance over time. The disaggregated molar and premolar data partitions do not contain large sample sizes; thus, we caution against over-interpreting the finer patterns until a statistical more robust sample can be analyzed. The per-tooth position analyses, although broadly supporting the early-middle Paleocene trends for both DTA and FEA data (and for FEA data in general across all time intervals), provide lower support for the middle-late Paleocene trends. Support is lowest when RFI and Slope values are analyzed by tooth position. In addition to the smaller sample sizes included in these partitioned analyses, this result may indicate that different tooth positions may record somewhat different, sometimes opposing, DTA signals but consistent FEA signals. This suggests structure-performance decoupling at the tooth position level. Similar differences between the pooled sample results and per-tooth trends when using range length (maximum trait value – minimum trait value for a given data partition) are recovered (Fig. S5B, 5D).

(3) Trait correlation patterns (Figs. S6-S7): To more precisely examine the relationships between pairs of DTA and FEA traits, we performed pairwise linear regression analyses using bootstrapped sample estimates of trait values. The outputs show that the 2B-PLS results below are mainly driven by association between higher DNE and OPCR with lower compressive and shear SE (Fig. S5). This association is particularly strong in premolar partitions and less strong in molar partitions (Fig. S6), and doesn't appear to represent differences between CEPs and non-pantodonts overall. When examined through time, there does seem to be a difference in how strongly DTA-FEA are associated; the spikes in Compressive and Shear SE values in the middle Paleocene time bin is not mirrored in DNE and OPCR values in the same time interval, suggesting a breakdown during middle Paleocene of the DTA-FEA relationship established in the early Paleocene. This result specifically mirrors findings from the CEP all-teeth data partition and non-pantodont molar partition 2B-PLS analysis (see below).

(4) Two-block partial least squares (2B-PLS) patterns (Table S1): The overall dataset exhibited a steady increase in *r* coefficient values from the early to the late Paleocene, indicating strengthening DTA-FEA integration over time. The CEP data partition showed a dip in integration in the middle Paleocene in the all-teeth partition, a steady decrease for premolar partitions, but the opposite pattern (a spike) in the molar partition. Non-pantodonts show a different pattern, where all-teeth and premolar partitions spike in integration during the middle Paleocene, whereas the molar partition dips in the middle Paleocene.

Sensitivity analyses of tooth size trends

Tooth disparity was highest during the early Paleocene time interval of our dataset; this is consistent both using variance and range length as the disparity measure (Fig. S8). The overall decrease in size disparity and variance across the Paleocene based on the total dataset was additionally tested in individual tooth position partitions (Table S3). The overall disparity trends are also observed in premolar and upper molar data partitions, whereas overall tooth size trends are observed mainly in the lower premolar 4 data partition. Decrease in mean tooth size is most consistently observed across multiple tooth partitions for the early Paleocene to late Paleocene comparison (and less clearcut for early-middle and middle-late Paleocene comparisons, respectively). Shifts during the middle Paleocene are variably supported depending on tooth position analyzed. Given that the middle Paleocene is represented by the smallest sample size, the uncertainty surrounding per-tooth position trends during this time period may be explained by the low sample available for this study.

Sample and methodological limitations

The highly fragmentary nature of early Cenozoic mammal fossils in Asia means that even the best preserved faunas studied herein contain substantial missing information. First, the absence of a high-resolution chronological framework prevents the fossil data from being analyzed on a continuous time axis; the binning of the samples into three main intervals within a 10-million-year period hinders additional hypotheses about the environmental and climatic correlations of the dental structure-performance results presented. Second, the uneven sampling of the available mammalian assemblage throughout the Paleocene sites in China limits the breadth of ecomorphological categories included in the analyses; rarer taxa representing potentially more specialized carnivore, insectivore, or herbivore forms were not included in our sampling. Third, the spatial discontinuity of stratigraphically younger (Eocene) and older (Cretaceous) mammal assemblages means that body size and ecomorphological shifts bracketing the Paleocene cannot currently be analyzed across the sampled basins alongside the dataset presented. These limitations should be considered when interpreting the findings reported in the study.

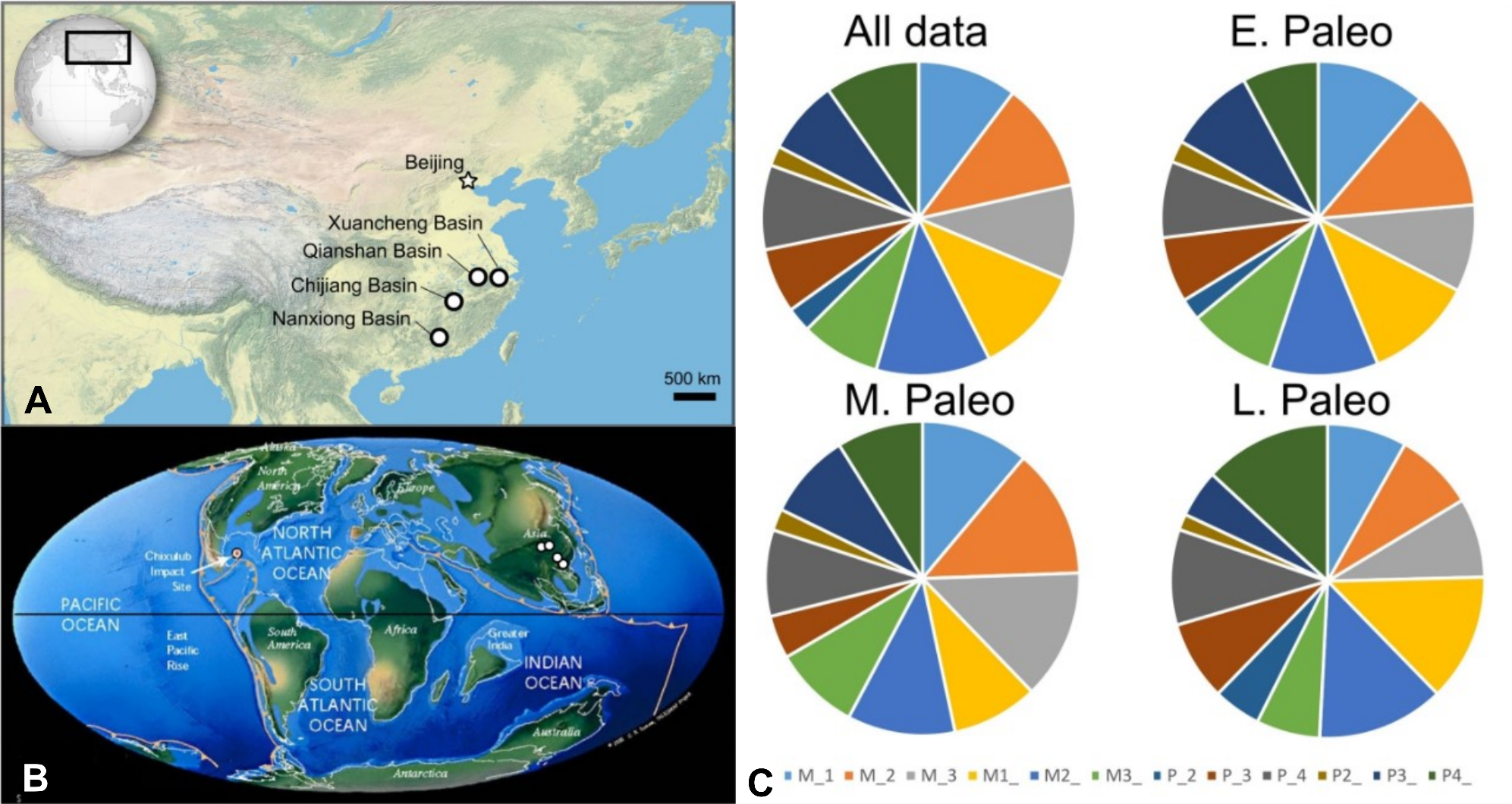

**Figure S1.** Maps of the fossil locality areas and proportions of teeth analyzed, related to Fig. 1. The Paleocene sedimentary basins sampled are indicated on a modern satellite image (**A**) and on a paleogeographic map of the K-Pg transition (**B**). Satellite map from NASA (www.nasa.gov) and paleogeographic map from C. Scotese (www.scotese.com)[20]. **C.** Proportions of tooth positions present in data partitions. M_1-M_3, lower first to third molars; M1_-M3_, upper first to third molars; P_2-P_4, lower second to fourth premolars; P2_-P4_, upper second to further premolars; E, early, M, middle, L, late. The middle Paleocene partition has no P_2 sampled, but otherwise the proportions of tooth positions present in each time interval are generally similar to each other.

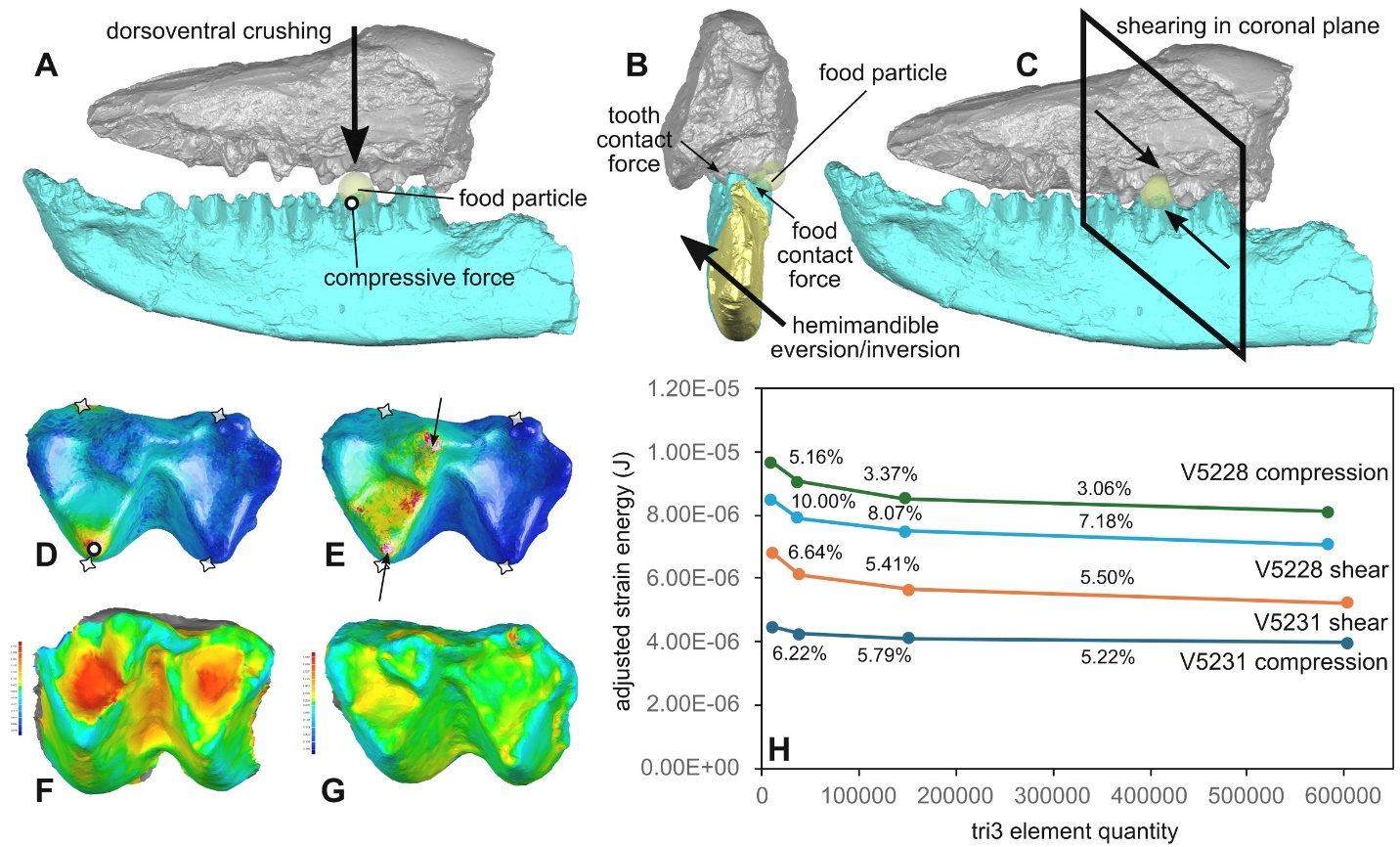

**Figure S2.** Finite element modeling boundary conditions and convergence tests, related to Figs. 1 and 3. (**A**) Depiction of compressive/crushing force imposed on a lower molar cusp in a specimen of *Guichilambda zhaii* (IVPP V12037) in lateral view. (**B**) Depiction of shearing motion during hemimandible eversion/inversion cycle in coronal view. (**C**) Depiction of shearing motion in lateral view. (**D**) Boundary conditions of compressive bite simulations on the lower first molar of *Altilambda tenuis* (IVPP V5231). (**E**) Boundary conditions of shearing bite simulations on *A. tenuis*. Nodal constraints are indicated by stars, compressive force by circle, and shearing forces by black arrows. (**F**) Three-dimensional deviation map of the first lower molar of *Altilambda pactus* (IVPP V5228) comparing geometric difference between models built form original versus cast-derived image data; positive (red) and negative (blue) deviation maxima +/- 0.58 mm. (**G**) Three-dimensional deviation map of the first lower molar of *Altilambda tenuis* (IVPP V5231), positive (red) and negative (blue) deviation maxima +/- 0.39 mm. (**H**) Convergence test of adjusted strain energy values (in Joules) of compression and shear bite simulations in IVPP V5228 and V5231. Percentage values indicate differences between adjacent model quantities (measured in number of three-noded triangular [tri3] elements). All model values at the lowest quantity models are considered converged based on a <=10% threshold criterion.

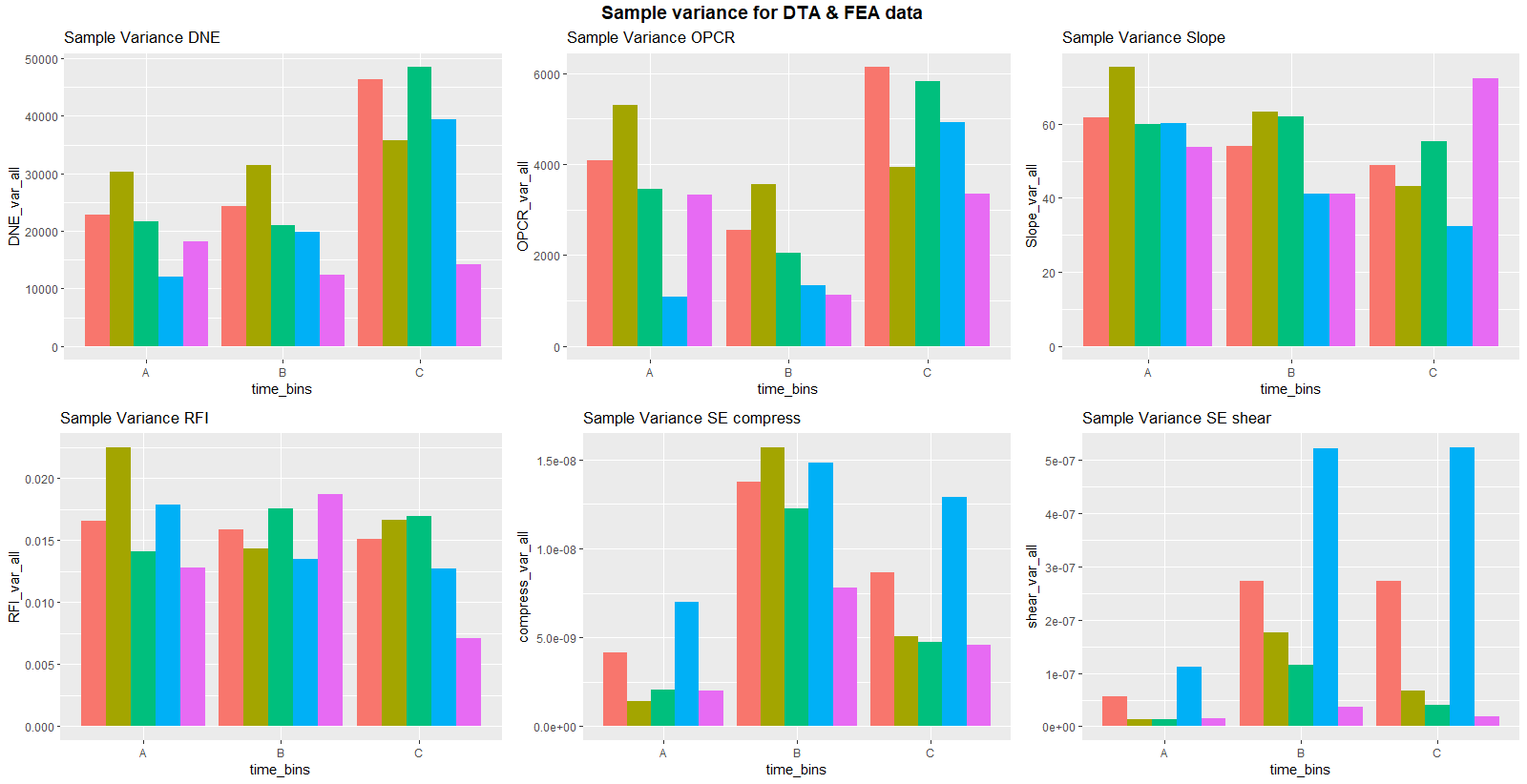

Figure S3. Sample variance (in squared units of each metric) of dental topographic metrics and dental performance traits for the overall dataset, related to Fig. 1. Time bins refer to the early Paleocene (Bin A), middle Paleocene (Bin B), and late Paleocene (Bin C). Color bars represent different data partitions: all data (salmon), all molars (teal), all premolars (green), lower molars (blue), and upper molars (purple).

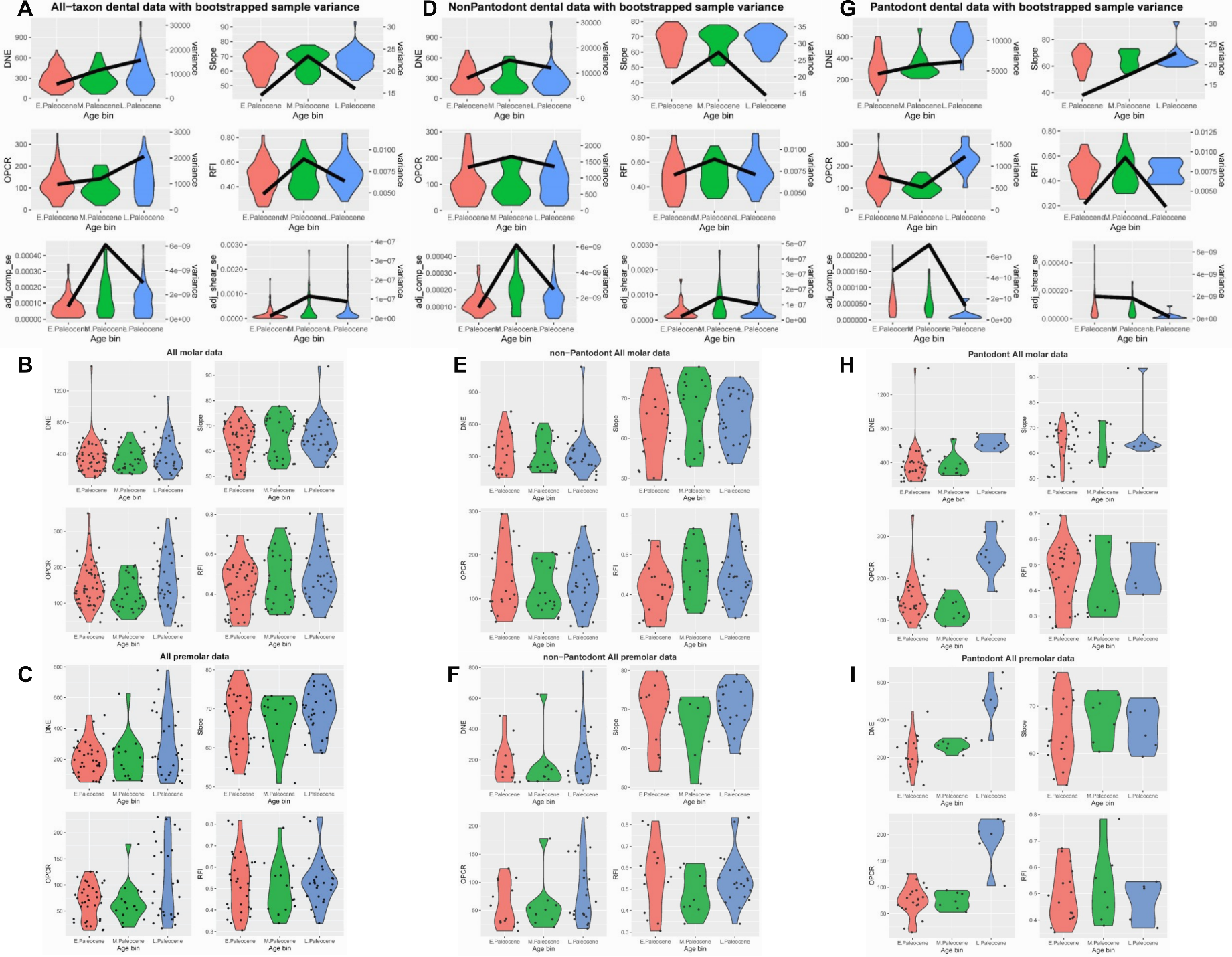

**Figure S4.** Boxplots of dental topographic and finite element simulated traits, related to Table 1. Dark lines in panels A, D, and G denote sample variance per million years, calculated from 1,000 bootstrap samples of trait values pulled from uniform distributions within their estimated uncertainty ranges. The variance magnitudes are scaled to the duration of the Asian land mammal ages representing each time interval. (**A**) DTA (DNE, OPCR, Slope, RFI) and FEA (compressive bite strain energy, shear bite strain energy) boxplots representing the all tooth positions in the all-taxon partition (**B**) DTA metrics for the molar partition of all taxa. (**C**) DTA metrics for the premolar partition of all taxa. (**D**) DTA and FEA boxplots of all tooth position in the non-pantodont data partition. (**E**) DTA metrics for molars in the non-pantodont data partition. (**F**) DTA metrics for premolars in the non-pantodont data partition. (**G**) DTA and FEA boxplots of all tooth positions in the Chinese endemic pantodont (CEP) partition. (**H**) DTA metrics for molars in the CEP partition. (**I**) DTA metrics for premolars in the CEP partition.

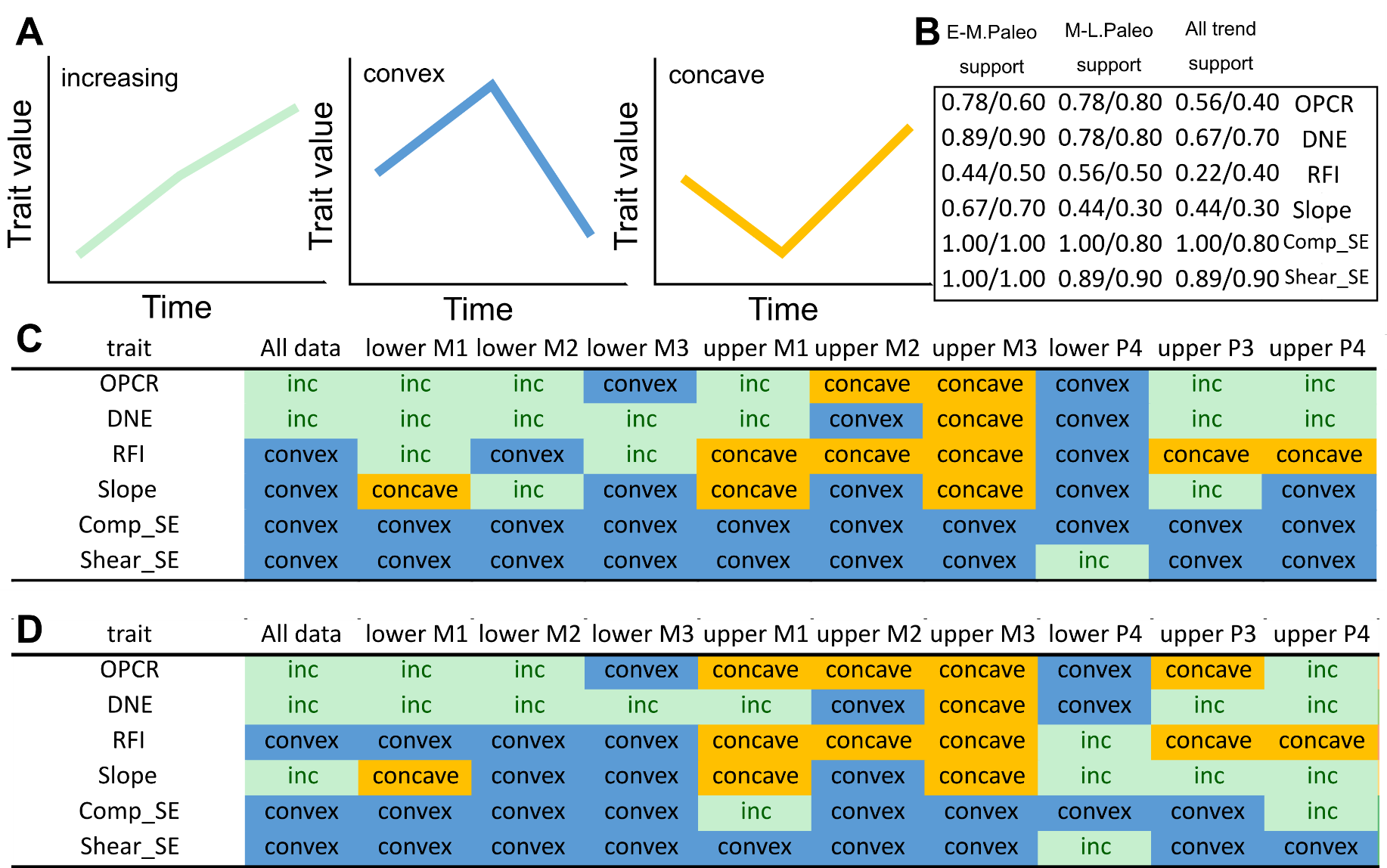

**Figure S5.** Trends through time for DTA and FEA traits by tooth position, related to Fig. 1. (**A**) Categories of major trends observed through the three Paleocene time intervals. (**B**) Proportion of individual tooth position analyses that support the trend observed in the overall dataset. The first set of values are per-tooth variance; the second set of values are from pooled sample and per-tooth range lengths. (**C**) Individual tooth position variance trends in DTA and FEA trait values. (**D**) Individual tooth position range length trends in DTA and FEA trait values.

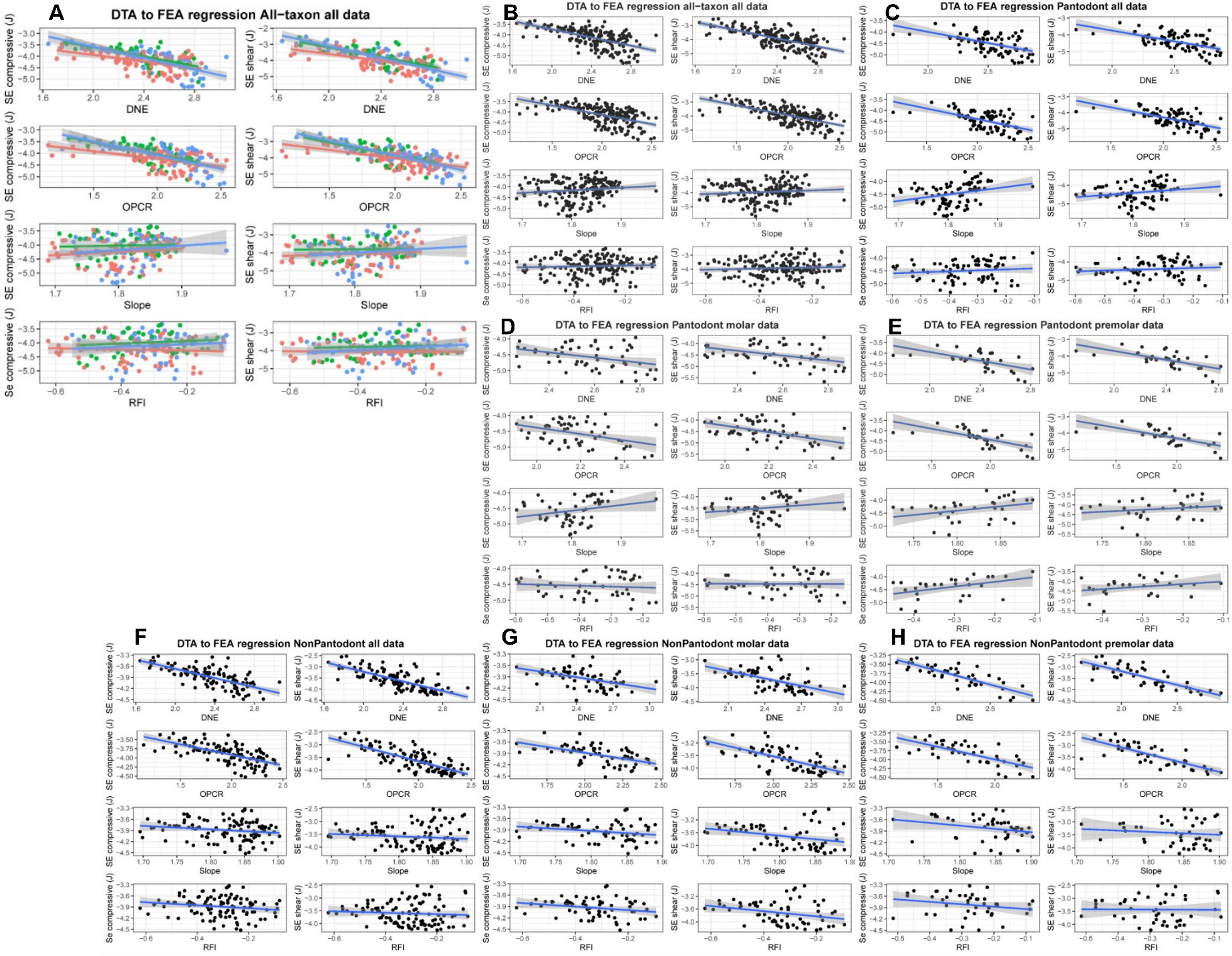

Figure S6. Linear regression analysis of dental topographic and bite performance datasets different data partitions, related to Fig. 3. (A) All-taxon all-teeth data partition; data points are colored according to early (red), middle (green), and late (blue) Paleocene age of the specimens. (B) All-taxon all-teeth data partition without time bine groups. (C) Chinese endemic pantodont (CEP) all-teeth data partition. (D) CPE molar data partition. (E) CEP premolar data partition. (F) Non-pantodont all-teeth partition. (G) Non-pantodont molar partition. (H) Non-pantodont premolar partition. Performance variables estimated using finite element analysis includes tooth crown strain energy under compressive (left column within each panel) and shear bite (right column within each panel) simulations, respectively. Rows within panels represent different DTA metrics (from top to bottom: DNE, OPCR, Slope, RFI). Fitted regression lines are shown in blue; 95% confidence envelopes are shown in gray shades.

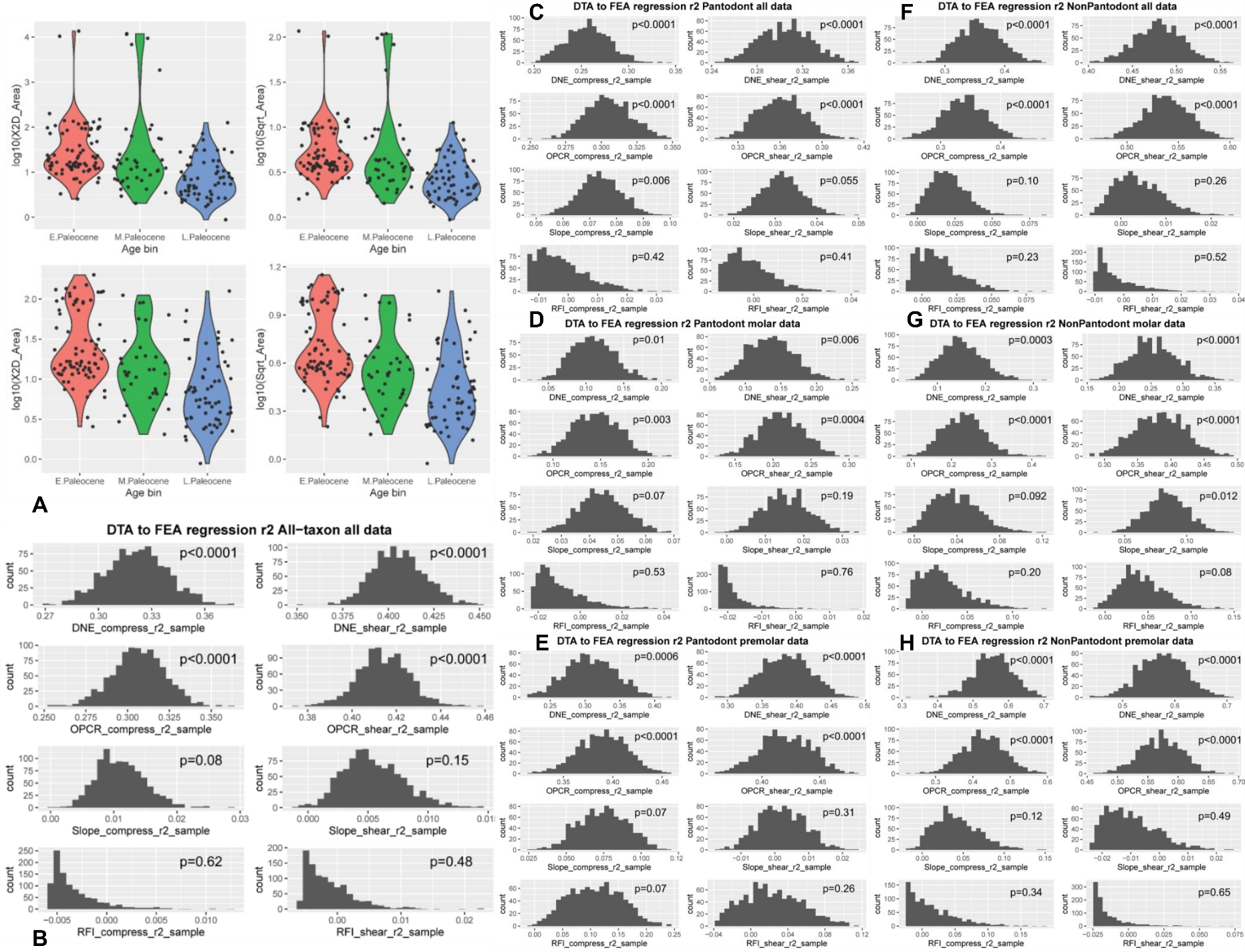

Figure S7. Tooth size boxplots and adjusted *r^2^* distributions from DTA-FEA linear regression analyses, related to Fig. 3. (A) Boxplots of tooth size by 2D area (left column within panel) or square root of 2D area (right column within panel). The top row within panel A includes sampling outliers (very large Chinese endemic pantodonts, CEPs), bottom row excludes outliers. The larger CEPs are removed in the bottom plots in order to more fully display the distribution of tooth sizes across the bimodal distribution of sizes in the main data cluster in each time bin, respectively.(B) Distribution of adjusted *r^2^* values from linear regression analysis of dental topographic and dental performance datasets for all-taxon all-teeth data partition. (C) CEP all-teeth partition. (D) CEP molar partition. (E) CEP premolar partition. (F) Non-pantodont all-teeth partition. (G) Non-pantodont molar partition. (H) Non-pantodont premolar partition. Adjusted *r^2^* values were generated from 1,000 bootstrap samples of DNE and FEA traits values for each specimen from uniform distributions that incorporate uncertainty in DNE and FEA values estimates. *P*-values calculated from *t* test of bootstrap sample *p* values against a hypothesis of *p* >= 0.05.

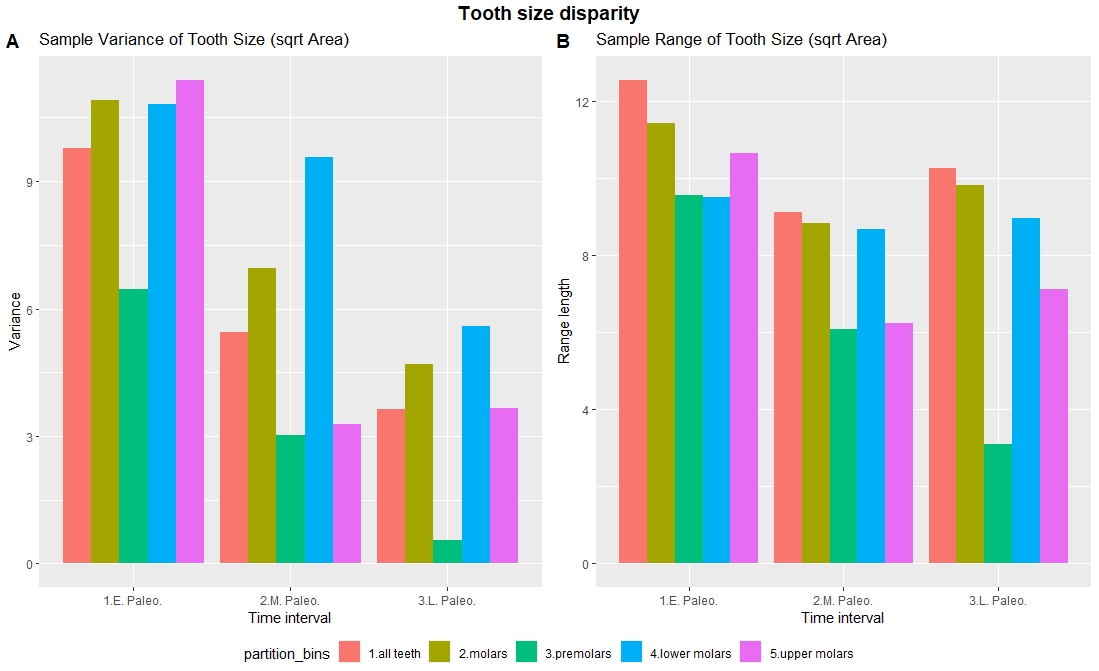

**Figure S8.** Tooth size disparity bar plots, related to Fig. 1. (**A**) Variance of tooth size (sqrt of 2D tooth area) in different data partitions. (**B**) Range length of tooth size (sqrt of 2D tooth area) in different partitions.

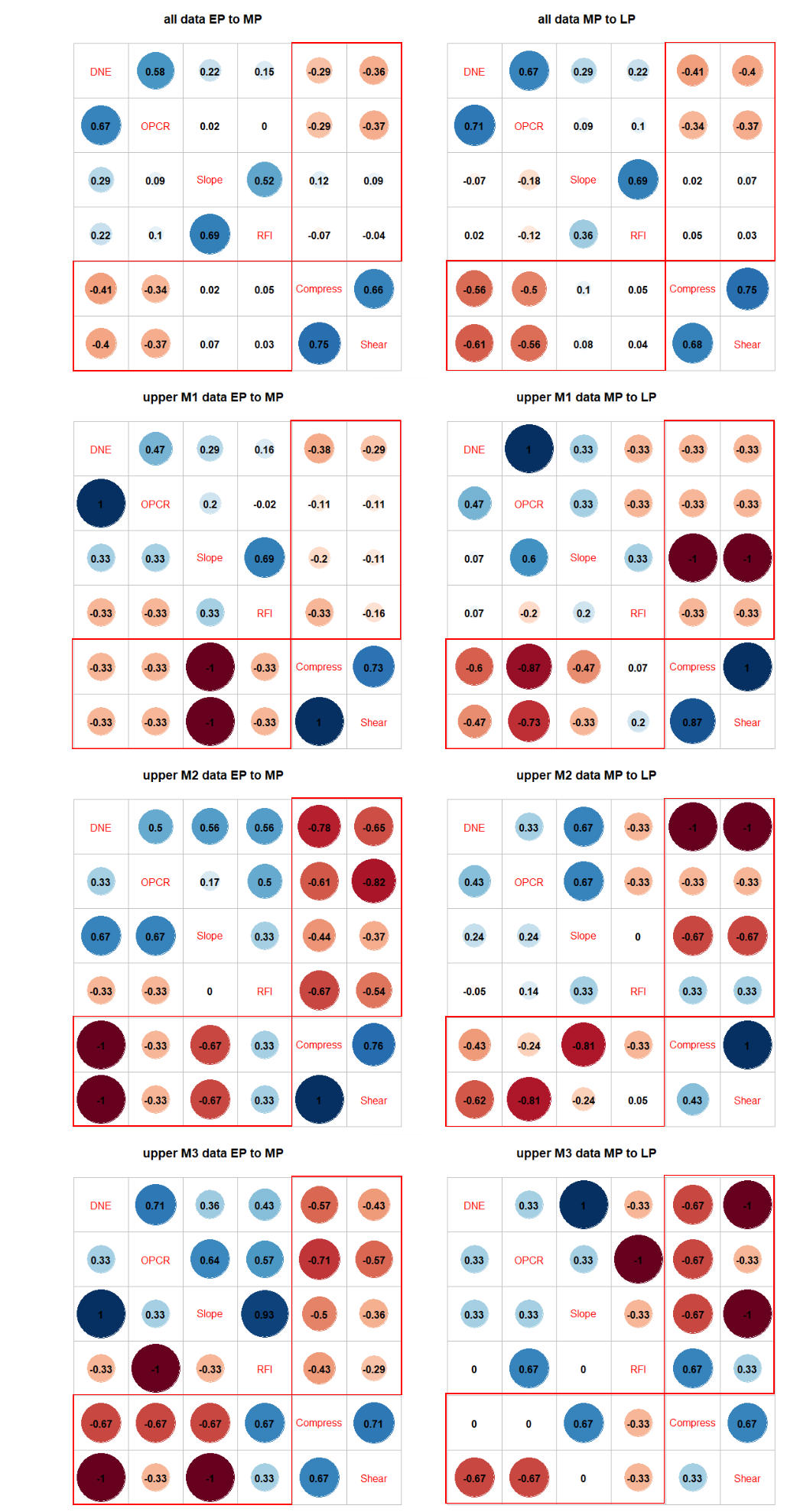

**Figure S9.** Correlation through time plots for upper molar DTA and FEA data, related to Fig. 3. The upper diagonal in the left column shows early Paleocene (EP) correlations, the lower diagonal in the left column shows middle Paleocene (MP) correlations. The upper diagonal in the right column shows middle Paleocene correlations, and the lower diagonal in the right column shows late Paleocene (LP) correlations. Red boxes indicate correlations between DTA and FEA traits. Dark gray boxes indicate decoupling of correlation directions across time intervals. Upper M1 patterns general reflect the trend recovered from analysis of the overall dataset, but M2 and M3 results display inconsistent DTA-FEA correlations, possibly due to small sample sizes.

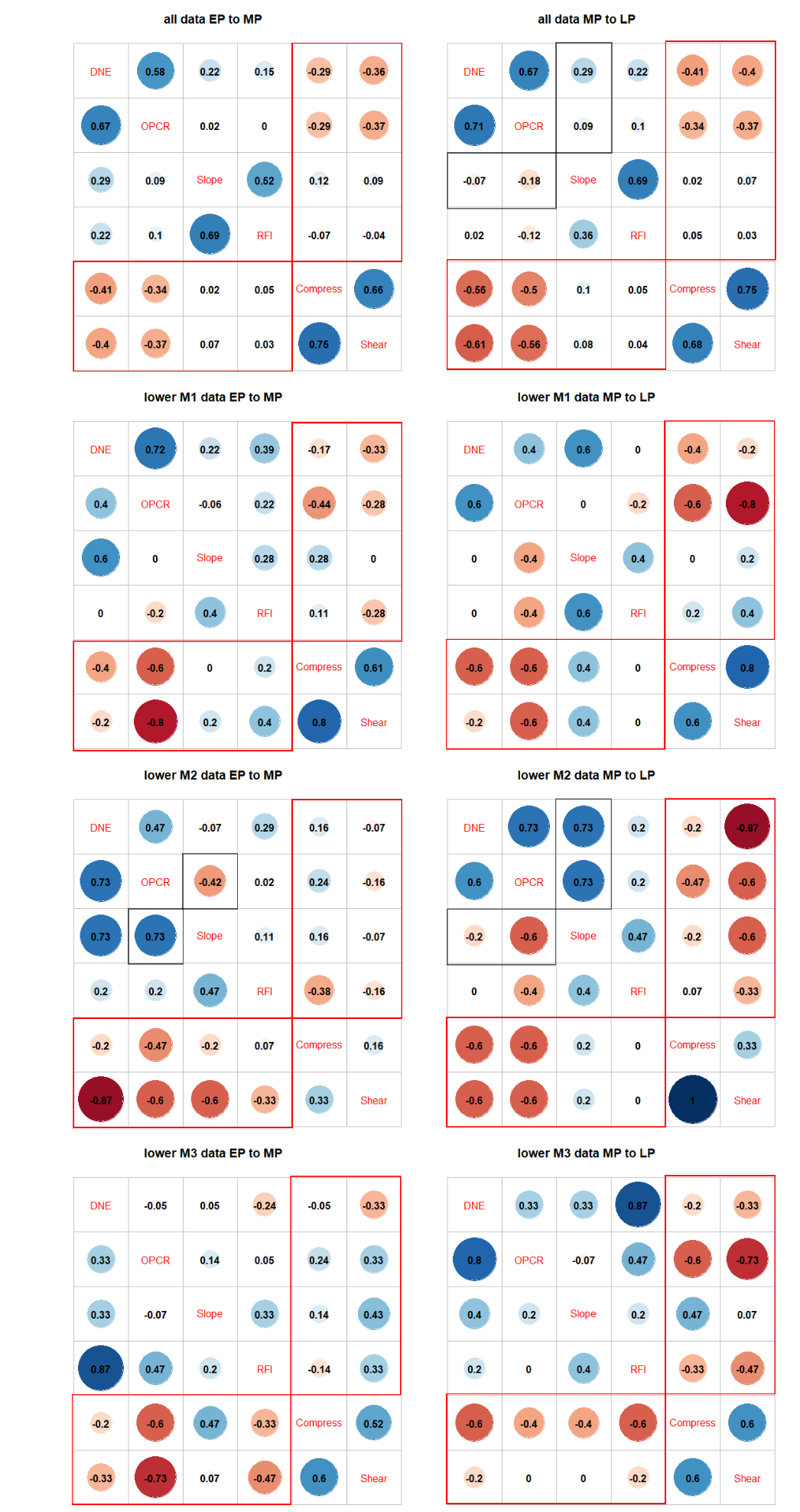

**Figure S10.** Correlation through time plots for lower molar DTA and FEA data, related to Fig. 3. The upper diagonal in the left column shows early Paleocene (EP) correlations, the lower diagonal in the left column shows middle Paleocene (MP) correlations. The upper diagonal in the right column shows middle Paleocene correlations, and the lower diagonal in the right column shows late Paleocene (LP) correlations. Red boxes indicate correlations between DTA and FEA traits. Dark gray boxes indicate decoupling of correlation directions across time intervals. Lower molar patterns generally replicate those recovered in the overall analyses, but lower M1 and M2 signals appear to be stronger than those for lower M3.

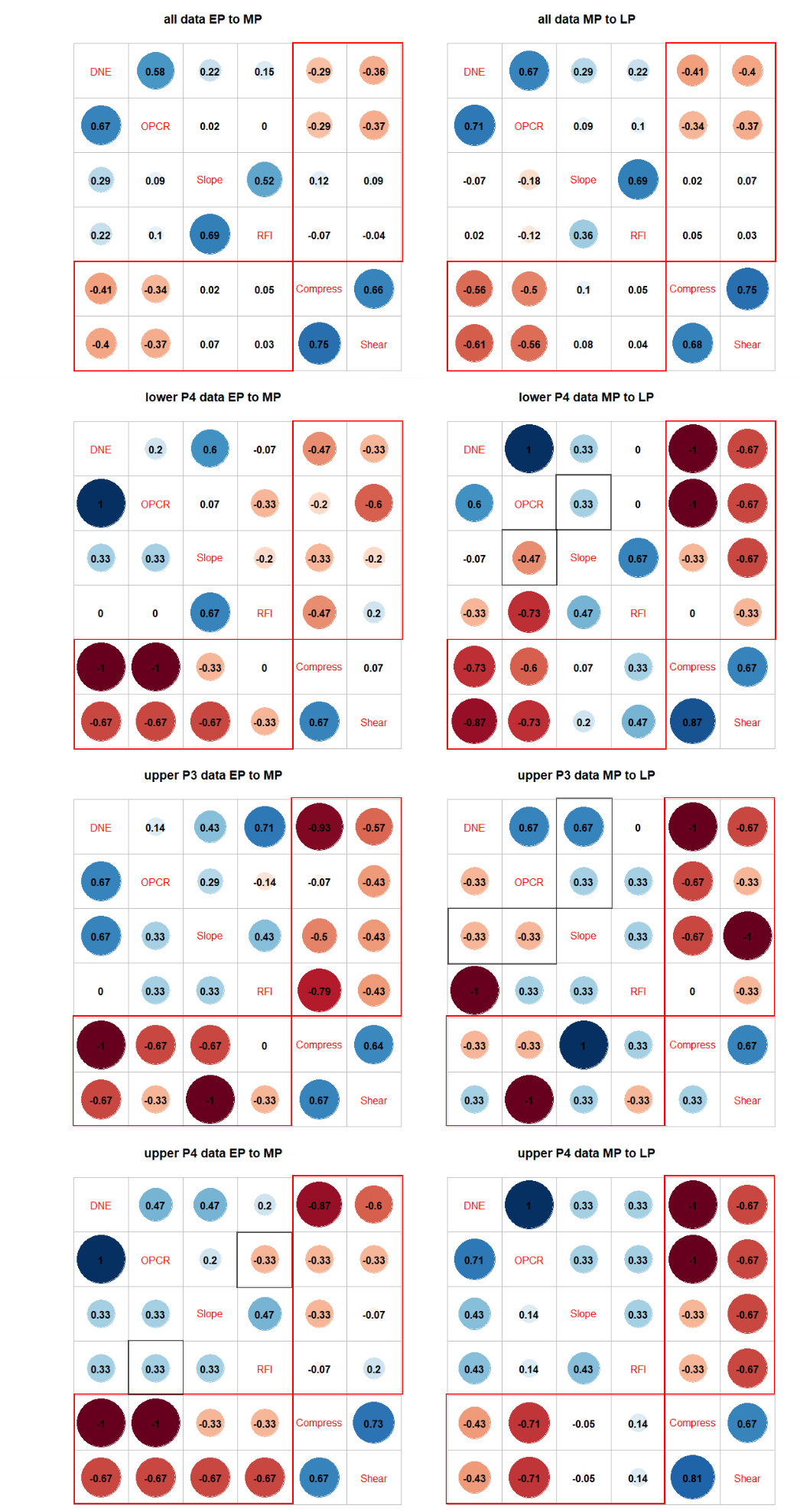

**Figure S11.** Correlation through time plots for premolar DTA and FEA data, related to Fig. 3. The upper diagonal in the left column shows early Paleocene (EP) correlations, the lower diagonal in the left column shows middle Paleocene (MP) correlations. The upper diagonal in the right column shows middle Paleocene correlations, and the lower diagonal in the right column shows late Paleocene (LP) correlations. Red boxes indicate correlations between DTA and FEA traits. Dark gray boxes indicate decoupling of correlation directions across time intervals. Low sample sizes make premolar correlations unstable, with general pattern showing EP-MP strengthening then MP-LP stasis or weakening

| Taxon | Dentition | early Paleocene | middle Paleocene | late Paleocene |
| --- | --- | --- | --- | --- |
| 1. All Taxa | All | 0.40 | 0.48 | 0.52 |
| 2. Chinese endemic pantodonts | All | 0.30 | 0.29 | 0.43 |
|  | Premolar | 0.69 | 0.57 | 0.51 |
|  | Molar | 0.15 | 0.70 | 0.65 |
| 3. Non-pantodont mammals | All | 0.48 | 0.53 | 0.49 |
|  | Premolar | 0.50 | 0.52 | 0.55 |
|  | Molar | 0.45 | 0.40 | 0.46 |

**Table S1.** Two-block partial least squares (PLS) *r* coefficients from bootstrapped analyses, related to Fig. 3. 1,000 bootstrap samples of DTA and FEA values were taken from uniform distributions of trait uncertainty ranges, and two-block partial east squared analysis conducted on each sample for early-middle Paleocene and middle-late Paleocene data partition pairings, respectively. All *r* values are statistically significantly different (*p* < 0.05) between adjacent time intervals, based on a one-sample t test of the distribution of 1,000 p values from the bootstrap two-block PLS samples against *p* < 0.05.

|  | E. Paleocene | | M. Paleocene | | L. Paleocene |
| --- | --- | --- | --- | --- | --- |
| Angiosperms | 75-88% | 84% | | 50-65% | |
| Ferns | 11-20% | 11% | | 25-33% | |
| Gymnosperms | 1-5% | 5% | | 10-18% | |

**Table S2.** Relative percentages of fossil pollen found in the Nanxiong Basin, Related to Fig. 2 and based on [3].

| **Measure** | **Trait** | **Statistic** | **All** | **m/1** | **m/2** | **m/3** | **M1/** | **M2/** | **M3/** | **p/4** | **P3/** | **P4/** |
| --- | --- | --- | --- | --- | --- | --- | --- | --- | --- | --- | --- | --- |
| Sample Size | Early Paleocene (EP) | | 79 | 10 | 11 | 8 | 10 | 10 | 8 | 7 | 8 | 7 |
|  | Middle Paleocene (MP) | | 42 | 5 | 6 | 6 | 4 | 5 | 4 | 4 | 4 | 4 |
|  | Late Paleocene (LP) | | 52 | 5 | 5 | 5 | 8 | 8 | 4 | 6 | 3 | 8 |
| Mean Disparity | Area | var.test EP-MP | 0.00 | 0.44 | 0.81 | 0.78 | 0.02 | 0.97 | 0.01 | 0.00 | 0.54 | 0.04 |
|  |  | var.test MP-LP | 0.03 | 0.00 | 0.06 | 0.92 | 0.14 | 0.00 | 0.04 | 0.08 | 0.05 | 0.24 |
|  | Sqrt Area | var.test EP-MP | 0.06 | 0.59 | 0.56 | 0.59 | 0.13 | 0.74 | 0.14 | 0.04 | 0.44 | 0.12 |
|  |  | var.test MP-LP | 0.16 | 0.02 | 0.11 | 0.72 | 0.28 | 0.06 | 0.07 | 0.30 | 0.17 | 0.72 |
| Mean Size | Area | t.test EP-MP | 0.00 | 0.18 | 1.00 | 1.00 | 0.75 | 0.91 | 0.06 | 0.04 | 0.86 | 0.53 |
|  |  | t.test MP-LP | 0.01 | 0.18 | 1.00 | 1.00 | 0.75 | 0.18 | 0.40 | 0.04 | 0.34 | 0.03 |
|  |  | t.test EP-LP | 0.00 | 0.01 | 1.00 | 1.00 | 0.75 | 0.02 | 0.01 | 0.00 | 0.34 | 0.00 |
|  | Sqrt Area | t.test EP-MP | 0.00 | 0.18 | 1.00 | 1.00 | 0.75 | 0.91 | 0.06 | 0.04 | 0.86 | 0.53 |
|  |  | t.test MP-LP | 0.01 | 0.18 | 1.00 | 1.00 | 0.75 | 0.18 | 0.40 | 0.04 | 0.34 | 0.03 |
|  |  | t.test EP-LP | 0.00 | 0.01 | 1.00 | 1.00 | 0.75 | 0.02 | 0.01 | 0.00 | 0.34 | 0.00 |

**Table S3**. Sample size, disparity, and mean tooth size by tooth position. Related to Fig. 1. Mean disparity difference is measured by pair-wise variance tests (var.test); mean tooth size difference is measured by pairwise *t* tests (t.test). *p* values <=0.05 are shaded. Overall disparity trends are also observed in premolar and upper molar data partitions, whereas overall tooth size trends are observed mainly in the lower premolar 4 data partition. Decrease in mean tooth size is most consistently observe across multiple tooth partitions for the early Paleocene to late Paleocene comparison.
